# Maternal progesterone signaling establishes lifelong oral homeostasis

**DOI:** 10.64898/2026.08.10.743932

**Authors:** Mayu Yamano, Junki Miyamoto, Daiki Sasahara, Yuki Masujima, Takako Ikeda, Ryuji Ohue-Kitano, Akari Nishida, Shingo Kawai, Rina Kurokawa, Nozomu Kono, Munekage Yamaguchi, Shinsuke Inuki, Fumitaka Osakada, Kentaro Nagaoka, Hiroaki Ohno, Yuki Sugiura, Nobuo Sasaki, Wataru Suda, Eiji Kondoh, Junken Aoki, Koji Hase, Ikuo Kimura

**Affiliations:** Department of Molecular Endocrinology, Graduate School of Pharmaceutical Sciences, Kyoto University, Sakyo-ku, Kyoto, Japan; Department of Applied Biological Science, Graduate School of Agriculture, Tokyo University of Agriculture and Technology, Fuchu, Tokyo, Japan; Laboratory of Molecular Metabolism, Graduate School of Biostudies, Kyoto University, Sakyo-ku, Kyoto, Japan; Laboratory of Biochemistry, Graduate School of Pharmaceutical Sciences, Ritsumeikan University, Kusatsu, Shiga, Japan; Department of Biological & Environmental Chemistry, Kindai University, Kayanomori, Iizuka, Fukuoka, Japan; Division of Biochemistry, Faculty of Pharmacy and Graduate School of Pharmaceutical Science, Keio University, Minato-ku, Tokyo, Japan; RIKEN Center for Integrative Medical Sciences, Yokohama, Kanagawa, Japan; Graduate School of Pharmaceutical Sciences, The University of Tokyo, Bunkyo-ku, Tokyo, Japan; Department of Obstetrics and Gynecology, Faculty of Life Sciences, Kumamoto University, Chuo-Ku, Kumamoto, Japan; Department of Bioorganic Medicinal Chemistry, Graduate School of Pharmaceutical Sciences, Kyoto University, Sakyo-ku, Kyoto, Japan; Laboratory of Cellular Pharmacology, Graduate School of Pharmaceutical Sciences, Nagoya University, Nagoya, Aichi, Japan; AMED-CREST, Japan Agency for Medical Research and Development, Chiyoda-ku, Tokyo, Japan; Laboratory of Veterinary Physiology, Graduate School of Agriculture, Tokyo University of Agriculture and Technology, Fuchu, Tokyo, Japan; Multi-Omics Platform, Center for Cancer Immunotherapy and Immunobiology, Graduate School of Medicine, Kyoto University, Sakyo-ku, Kyoto, Japan; Laboratory of Mucosal Ecosystem Design, Institute for Molecular and Cellular Regulation, Gunma University, Maebashi, Gunma, Japan; Department of Moonshot Research and Development Program, Japan Science and Technology Agency, Chiyoda-ku, Tokyo, Japan; Division of Commensal Biology, The Institute of Medical Science, The University of Tokyo (IMSUT), Bunkyo-ku, Tokyo, Japan; International Vaccine Design Center, The Institute of Medical Science, The University of Tokyo (IMSUT), Bunkyo-ku, Tokyo, Japan

**Keywords:** mPRδ, Paqr6, progesterone, membrane progesterone receptor

## Abstract

Pregnancy is accompanied by profound endocrine remodeling, yet the mechanisms by which maternal hormonal signals establish long-term tissue homeostasis remain largely unknown. Here we identify maternal progesterone signaling as a developmental cue that establishes lifelong oral homeostasis through a hormone–lipid–microbiome axis. We show that the membrane progesterone receptor mPRδ is selectively expressed in the developing and maternal submandibular glands, where it mediates non-genomic progesterone signaling to promote epithelial differentiation by driving the selective mobilization of docosahexaenoic acid (DHA). Loss of this pathway disrupts salivary gland maturation, reshapes the oral microbial ecosystem through the selective expansion of Pasteurellaceae, and causes local inflammation as well as systemic metabolic dysfunction. Mechanistically, antibiotic treatment abolishes these phenotypes, whereas transfer of the oral microbiota recapitulates disease, demonstrating that developmental defects in the host are translated into long-term pathology through the oral microbiome. Remarkably, maternal—but not adult—DHA supplementation restores salivary gland development and microbial homeostasis and prevents adult disease phenotypes, identifying a critical developmental window during which oral homeostasis is durably established. Collectively, these findings reveal a previously unrecognized maternal endocrine mechanism that establishes lifelong host–microbiome homeostasis and identify developmental programming as a fundamental principle linking maternal physiology to adult health.

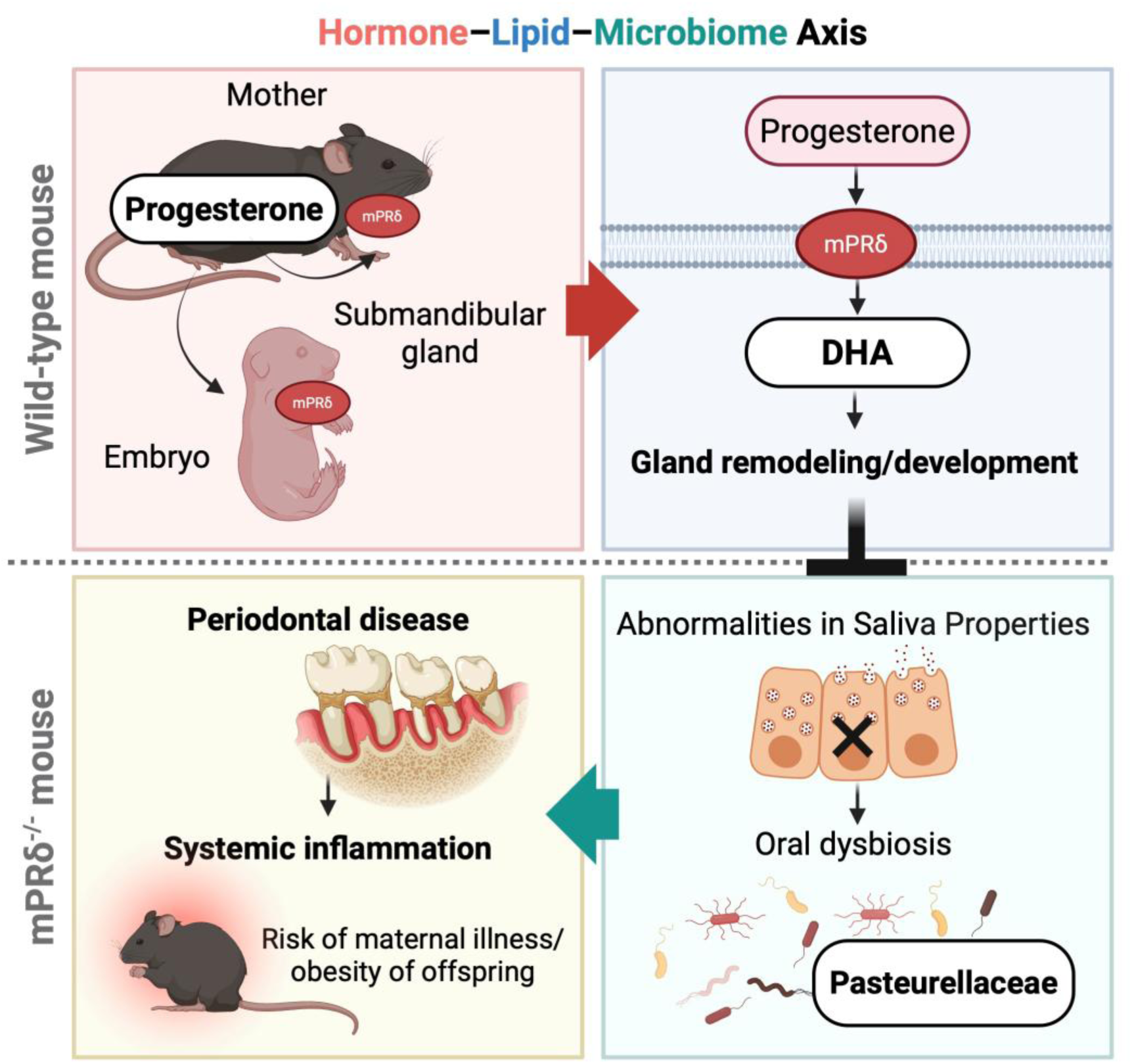

## Introduction

The oral cavity constitutes a major host–environment interface whose homeostasis influences physiology far beyond the oral cavity itself^1–4^. In addition to classical oral diseases such as dental caries and periodontal disease, alterations in salivary composition and oral microbial communities have been implicated in cardiovascular disease, metabolic disorders, adverse pregnancy outcomes, and neuroinflammatory conditions^1–4^. For example, periodontal disease is associated with insulin resistance and atherosclerosis, whereas pregnancy-associated oral inflammation correlates with preterm birth and low birth weight^5–7^. These observations highlight the importance of oral homeostasis in systemic health. However, despite pronounced sex differences in many oral diseases and the heightened susceptibility to oral inflammation during pregnancy^8–10^, the mechanisms by which maternal endocrine signals remodel the oral environment and establish long-term oral homeostasis remain largely unknown.

Saliva plays a central role in maintaining oral homeostasis by regulating antimicrobial defense, immune surveillance, and microbial ecology, with the submandibular gland serving as the principal source of functional salivary proteins^11,12^. The gland exhibits marked sexual dimorphism, exemplified by the development of the granular convoluted tubule (GCT) in rodents, and contains multiple epithelial lineages, including basal, intercalated duct (ID), and striated duct (SD) cells^11,13–15^. These specialized cell types arise through the coordinated differentiation of immature ductal progenitors during late fetal and early postnatal development^11,13–15^. These observations suggest that temporally restricted developmental cues establish glandular architecture and secretory function. However, the molecular mechanisms by which endocrine signals coordinate epithelial lineage specification and functional maturation during this critical developmental window remain poorly understood.

Maternal progesterone is a major endocrine signal that orchestrates physiological adaptation during pregnancy, increasing to concentrations more than an order of magnitude higher than those observed during the estrous cycle^16,17^. Pregnancy is accompanied by extensive remodeling of the salivary proteome^18^. However, many of these changes occur on a timescale inconsistent with transcriptional regulation mediated by classical nuclear progesterone receptors^19,20^. This suggests the involvement of rapid non-genomic signaling mechanisms. Membrane progesterone receptors (mPRs), members of the progestin and adipoQ receptor (PAQR) family, have emerged as mediators of rapid progesterone signaling^21–25^. These receptors activate diverse intracellular pathways, including MAPK signaling, calcium mobilization, and phospholipase-dependent lipid remodeling^21–25^. Although accumulating evidence from *in vitro* studies has implicated mPRδ in multiple cellular processes^23,26^, its physiological function *in vivo*, particularly during organ development and maternal endocrine adaptation, remains essentially unknown.

Here, building on our previous studies identifying maternal–offspring communication mediated by the gut microbiota^27^ and the membrane progesterone receptor ε (mPRε)^25^, we identify mPRδ as a developmental mediator of maternal progesterone signaling in the fetal and maternal submandibular glands. We demonstrate that maternal progesterone signaling directs salivary gland development to establish lifelong oral homeostasis through a hormone–lipid–microbiome axis. These findings uncover a developmental mechanism by which maternal endocrine signals program host–microbiome interactions and systemic physiology.

## Results

### Progesterone-mPRδ signaling regulates submandibular gland development

We first characterized progesterone dynamics and mPRδ expression during the perinatal period to determine whether progesterone regulates submandibular gland development through mPRδ. Maternal and fetal plasma progesterone concentrations increased to >300 nM and ∼30 nM, respectively, during pregnancy, levels sufficient to activate members of the mPR family^23^ (Fig. 1a). Among all mPR family members, only *mPR*δ was prominently expressed in the submandibular gland at embryonic day 18.5 (E18.5) (Fig. 1b). *mPR*δ mRNA expression peaked during the perinatal period in both male and female offspring (Fig. 1c). It remained relatively abundant in adult submandibular glands compared with that in other tissues, including the lung, liver, colon, and ovary (Extended Data Fig. 1a). In contrast, expression of the nuclear progesterone receptor (*Pgr*) was negligible in the submandibular gland relative to its expression in the ovary (Extended Data Fig. 1b).

**Fig. 1.**
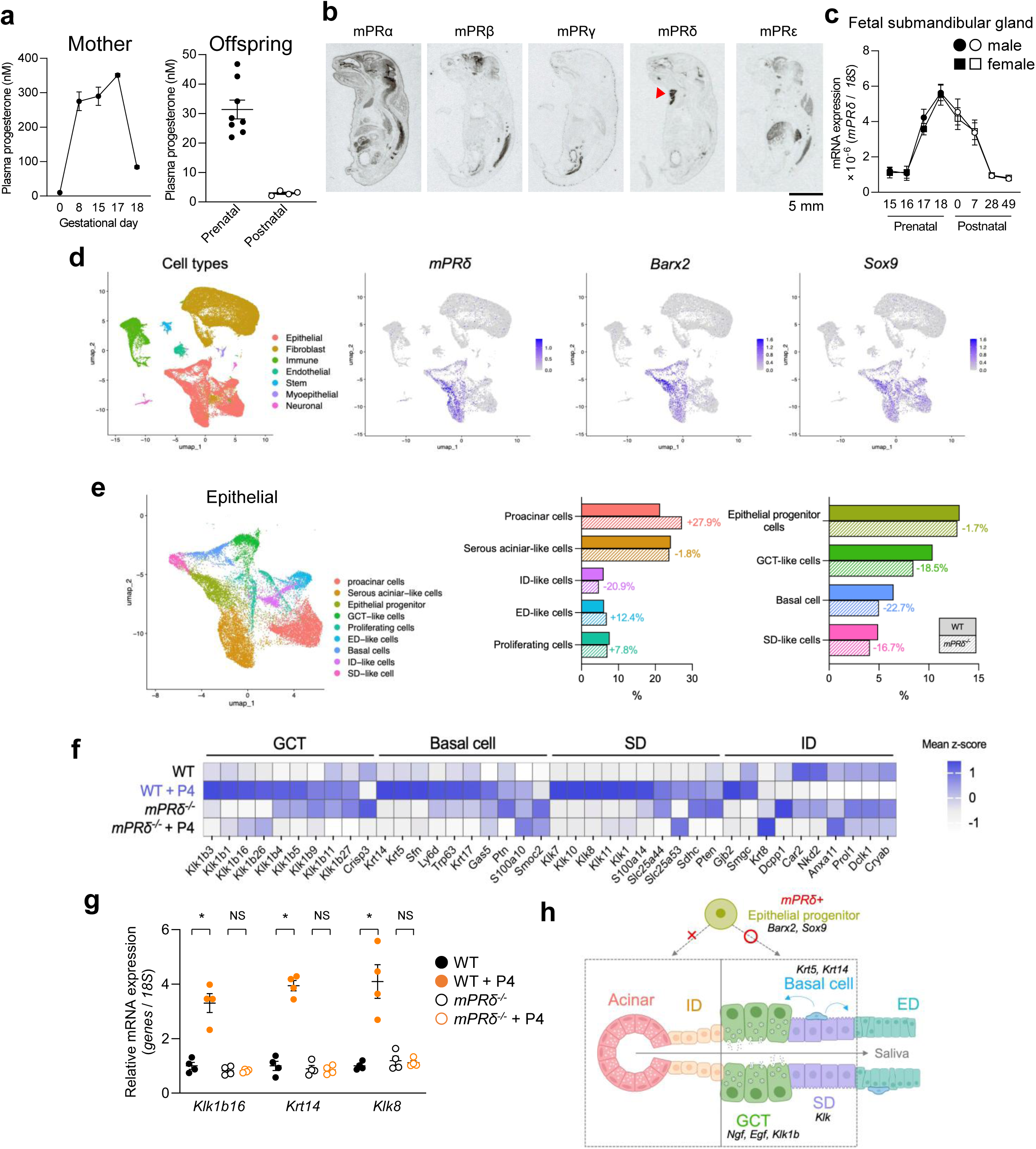
| Progesterone–mPRδ signaling regulates epithelial lineage specification in the developing submandibular gland. (a) Progesterone concentrations in maternal plasma across gestation (left) and in offspring plasma at prenatal and postnatal stages (right) (mother: *n* = 3–4; offspring: *n* = 4–8). (b) *In situ* hybridization showing expression patterns of mPR family members (*mPR*α, *mPR*β, *mPR*γ, *mPR*δ, and *mPR*ε) in mouse embryos at embryonic day 18.5 (E18.5). The arrowhead indicates prominent *mPR*δ expression in the submandibular glands (SMGs). Scale bar, 5 mm. (c) Temporal expression of *mPR*δ mRNA in fetal and postnatal SMGs from male and female mice (*n* = 3–5). (d) Uniform manifold approximation and projection (UMAP) visualization of single-cell RNA-seq data from SMGs at postnatal day 0 (P0), annotated by major cell types (left). Feature plots showing expression of *mPR*δ and the epithelial progenitor markers *Barx2* and *Sox9* (right). (e) UMAP visualization of epithelial cells extracted from the dataset in (d) (left). Bar plots showing the proportion of epithelial subpopulations in wild-type and *mPR*δ^□^*^/^*^□^ mice (right). (f) Heatmap of gene expression in SMG organoids derived from wild-type or *mPR*δ^□^*^/^*^□^ embryos treated with vehicle or progesterone (P4; 100 nM) (*n* = 3). (g) RT-qPCR analysis of epithelial differentiation markers in SMG organoids (*n* = 4). Expression was normalized to 18S rRNA. (h) Proposed model illustrating how progesterone–mPRδ signaling regulates epithelial progenitor differentiation and lineage specification in the developing SMGs. Data are presented as mean ± SEM. Statistical significance was determined using a two-tailed Student’s *t*-test or one-way ANOVA followed by the appropriate post hoc tests, as indicated (\**P* < 0.05; NS, not significant).

We performed single-cell RNA sequencing (scRNA-seq) of postnatal day 0 (P0) submandibular glands to define the cellular distribution of *mPR*δ. *mPR*δ expression was restricted to a subset of epithelial progenitor cells expressing *Barx2* and *Sox9*^28,29^ (Fig. 1d). In contrast, little or no expression was detected in basal, acinar, SD or GCT cells (Extended Data Fig. 1c, d). These findings indicate that mPRδ is preferentially expressed in a subset of epithelial progenitors during perinatal submandibular gland development and suggest a role in epithelial lineage specification.

We generated mPRδ-deficient mice (Extended Data Fig. 2a–c) to investigate the physiological function of mPRδ. Although these mice showed no overt developmental abnormalities other than mild hyperglycemia in both sexes (Extended Data Fig. 2d–f), scRNA-seq analysis of P0 submandibular glands revealed a selective reduction in differentiated epithelial populations. Whereas epithelial progenitor cells were preserved, GCT-like, SD-like, and basal-cell populations^30^ were markedly reduced in mPRδ-deficient mice (Fig. 1e and Extended Data Fig. 1e), indicating impaired epithelial differentiation.

We established primary organoid cultures from embryonic submandibular glands (Extended Data Fig. 3a, b) to determine whether progesterone directly promotes epithelial differentiation through mPRδ. RNA sequencing followed by Gene Ontology (GO) enrichment analysis revealed that progesterone induced transcriptional programs associated with epithelial differentiation and epidermal development in wild-type organoids. In contrast, these responses were largely absent in mPRδ-deficient organoids (Fig. 1f and Extended Data Fig. 3c, d). Consistent with these findings, progesterone significantly increased the expression of epithelial differentiation markers, including *Klk1b16*, *Krt14*, and *Klk8*^13–15^, in wild-type but not mPRδ-deficient organoids (Fig. 1g). Collectively, these findings establish progesterone–mPRδ signaling as a developmental regulator of epithelial lineage specification in the submandibular gland (Fig. 1h).

### mPR**δ**-dependent DHA mobilization drives epithelial differentiation in the submandibular gland

We next examined lipid remodeling in the developing submandibular gland to investigate the molecular mechanism by which mPRδ promotes epithelial differentiation. Previous studies have linked mPR signaling to phospholipase A2 (PLA2)-dependent lipid remodeling and fatty acid mobilization^24,25^. We therefore analyzed membrane phospholipid composition in P0 submandibular glands from mPRδ-deficient mice. Membrane phospholipid profiling revealed genotype-dependent alterations in several phospholipid species. Among these, PS 40:6 and ether-linked PE O-38:7 were relatively abundant and showed particularly pronounced accumulation in mPRδ-deficient mice (Fig. 2a and Extended Data Fig. 4a). Notably, both species are compatible with DHA-containing membrane phospholipids, suggesting impaired DHA mobilization from membrane phospholipid pools. Consistent with this interpretation, targeted profiling of non-esterified fatty acids revealed a marked reduction in free DHA in mPRδ-deficient submandibular glands (Fig. 2b, c). Although free eicosapentaenoic acid (EPA; 20:5) was also reduced, its abundance was more than an order of magnitude lower than that of DHA. Likewise, DHA-derived lipid mediators, including 16,17-epoxydocosapentaenoic acid (16,17-EpDPA), were detected at levels more than two orders of magnitude lower than free DHA, despite showing genotype-dependent changes (Fig. 2d and Extended Data Fig. 4b). Collectively, these findings pointed to DHA as the predominant free fatty acid regulated by mPRδ in the developing submandibular gland.

**Fig. 2.**
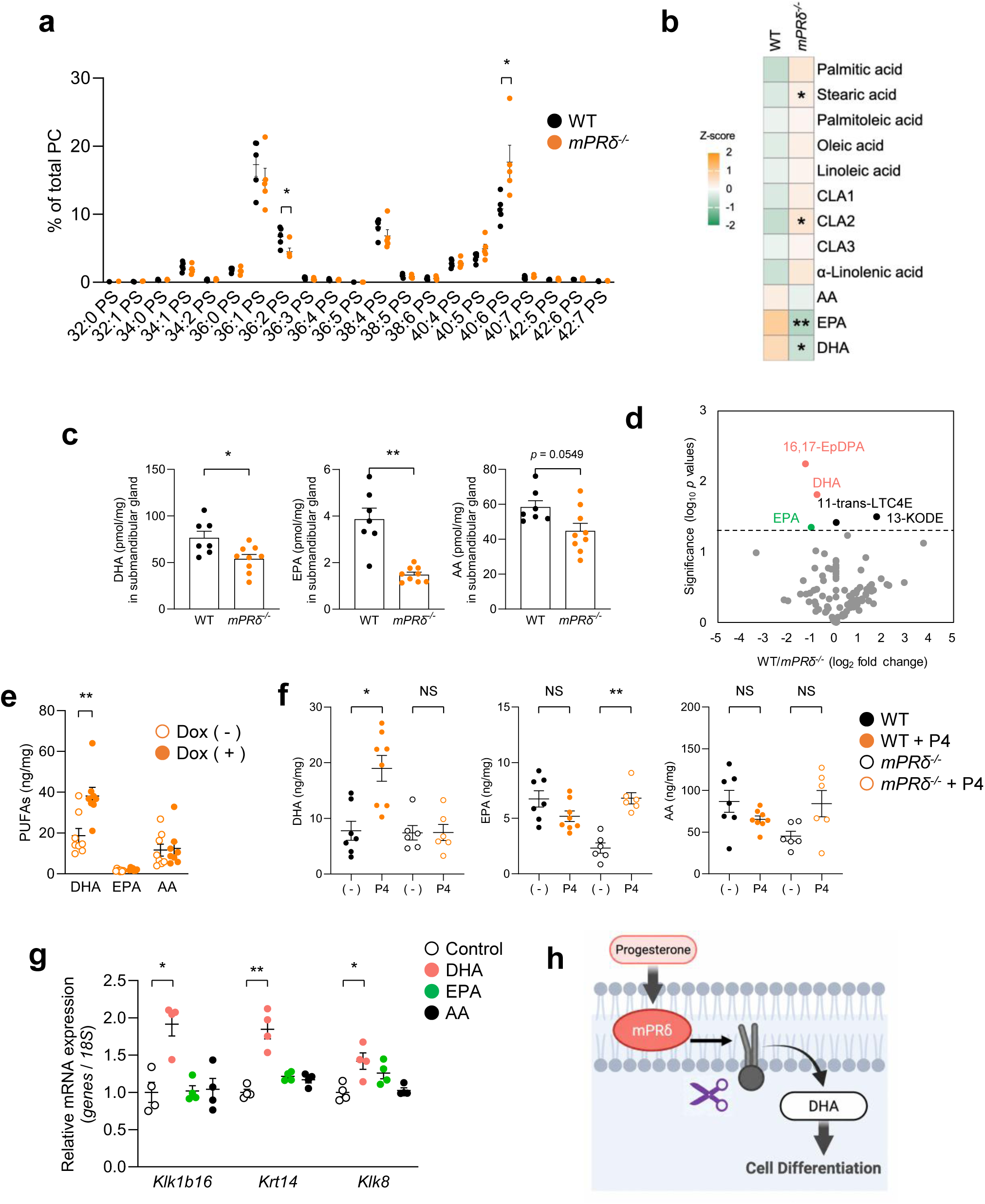
| mPRδ-dependent DHA mobilization promotes epithelial differentiation in the developing submandibular gland. (a) Quantification of phosphatidylserine (PS) species in SMGs from wild-type and *mPR*δ^□^*^/^*^□^ mice at P0 (*n* = 5). Values are expressed as a percentage of total phosphatidylcholine (PC) content. (b) Heatmap of the targeted profiling of non-esterified/free fatty acids of SMGs from wild-type and *mPR*δ^□^*^/^*^□^ mice at P0 (*n* = 6–10). Asterisks indicate significantly altered fatty acids (\**P* < 0.05, \*\**P* < 0.01). (c) Levels of free docosahexaenoic acid (DHA), eicosapentaenoic acid (EPA), and arachidonic acid (AA) in SMGs at P0 (*n* = 7–9). (d) Profiling of 193 lipid mediators derived from DHA, EPA, and AA in SMGs at P0 (*n* = 4). Differentially abundant lipid mediators are indicated. (e) Intracellular levels of free DHA, EPA, and AA in Flp-In T-REx HEK293 cells expressing mPRδ following progesterone (P4; 100 nM) stimulation. mPRδ expression was induced with doxycycline (Dox; 10 μg/mL) (*n* = 8). (f) Intracellular levels of free DHA, EPA, and AA in SMG–derived cells from wild-type and *mPR*δ^□^*^/^*^□^ embryos following treatment with progesterone (P4; 100 nM) (*n* = 6–8). (g) RT-qPCR analysis of epithelial differentiation markers in SMG organoids treated with DHA, EPA, or AA (*n* = 4). Expression was normalized to 18S rRNA. (h) Proposed model in which progesterone–mPRδ signaling promotes selective DHA mobilization from membrane phospholipids, thereby driving epithelial differentiation in the developing SMG. Data are presented as mean ± SEM. Statistical significance was determined using a two-tailed Mann–Whitney U test or one-way ANOVA followed by the appropriate post hoc tests, as indicated (\**P* < 0.05, \*\**P* < 0.01; NS, not significant).

To determine whether progesterone promotes DHA mobilization through mPRδ, we performed complementary *in vitro* analyses. In a heterologous expression system, progesterone did not activate canonical GPCR signaling pathways, including cAMP production or intracellular Ca² mobilization, in HEK293 cells expressing mouse mPRδ (Extended Data Fig. 5a–c). Under progesterone-stimulated conditions, mPRδ-expressing HEK293 cells exhibited selectively higher intracellular free DHA levels than control cells lacking mPRδ expression, whereas EPA and arachidonic acid (AA) levels were unchanged (Fig. 2e). Importantly, in primary submandibular gland organoids, progesterone stimulation increased intracellular free DHA levels in wild-type organoids but not in mPRδ-deficient organoids (Fig. 2f), establishing that progesterone-dependent DHA mobilization requires mPRδ.

We examined organoids derived from embryonic submandibular epithelial progenitors to determine whether DHA functions downstream of mPRδ. DHA, but not EPA or AA, robustly induced the expression of epithelial differentiation markers, including *Klk1b16*, *Krt14*, and *Klk8* (Fig. 2g). Collectively, these findings demonstrate that progesterone–mPRδ signaling promotes epithelial differentiation by selectively mobilizing DHA from membrane phospholipid pools, identifying DHA as the principal downstream effector of mPRδ during submandibular gland development (Fig. 2h).

### mPR**δ** deficiency disrupts salivary gland function and oral homeostasis in a sex-dependent manner

We examined salivary gland architecture and function in adult mPRδ-deficient mice to determine whether impaired epithelial differentiation results in functional defects *in vivo*. Histological analysis of 7-week-old males revealed marked abnormalities in ductal morphology, particularly within the SD, compared with that in wild-type controls (Fig. 3a). Consistent with these structural changes, principal component analysis (PCA) of transcriptomic profiles demonstrated a clear separation between wild-type and mPRδ-deficient submandibular glands in males. In contrast, glands in females remained largely unaffected (Fig. 3b). scRNA-seq analysis further revealed that sexual dimorphism in the adult submandibular gland is largely driven by the presence of a distinct GCT population (Extended Data Fig. 6a, b). Accordingly, mPRδ deficiency selectively reduced secretory GCT cells in males, whereas non-secretory SD populations were preferentially affected in females (Fig. 3c and Extended Data Fig. 6c, d). These findings indicate that mPRδ is required for maintaining epithelial lineages in a sex-dependent manner.

**Fig. 3.**
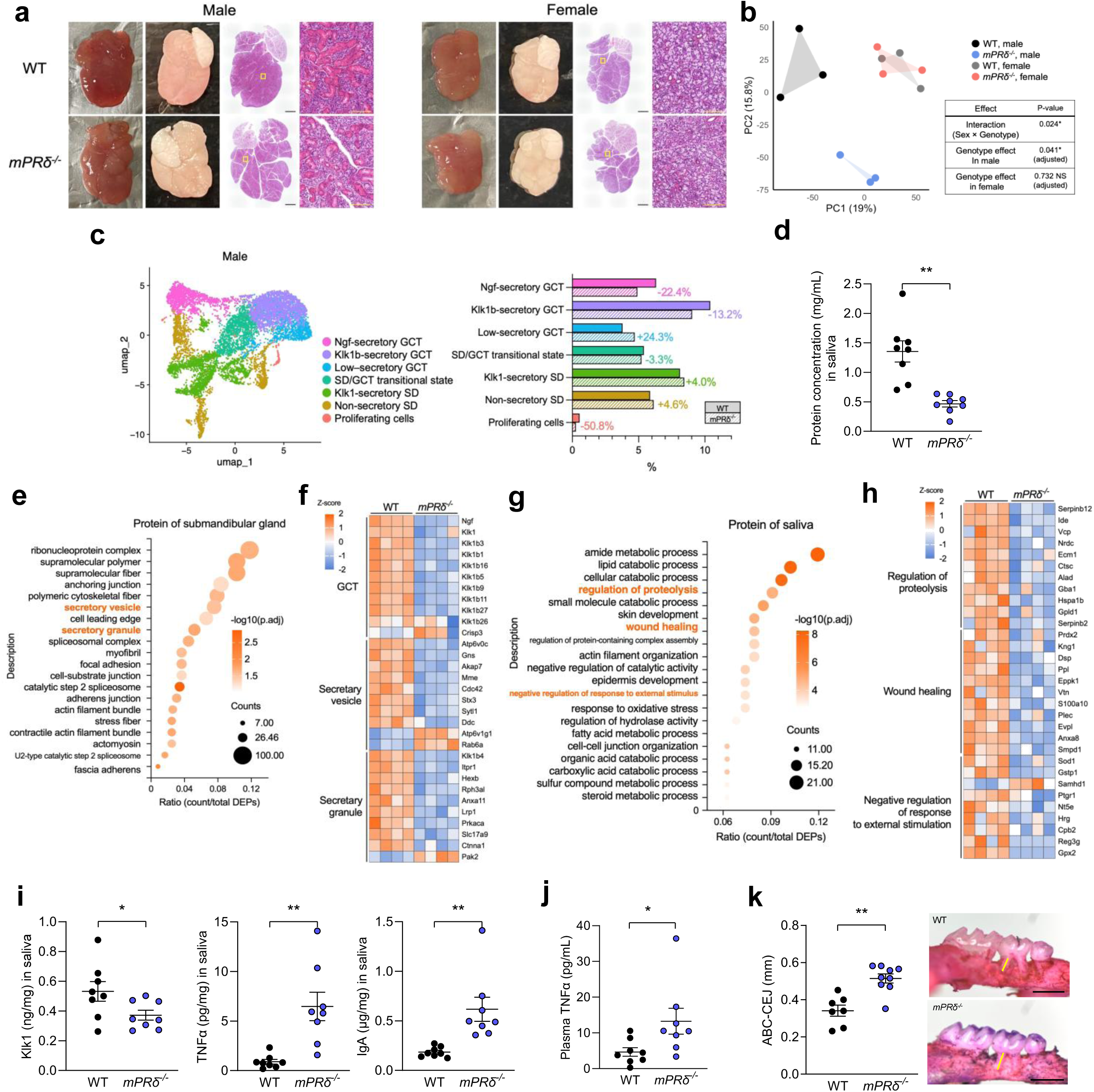
| Loss of mPRδ impairs salivary gland function and oral homeostasis in a sex-dependent manner. (a) Morphological and histological analysis of SMGs from 7-week-old male and female wild-type and *mPR*δ^□^*^/^*^□^ mice. Representative images show freshly dissected glands, dehydrated tissues and hematoxylin and eosin (H&E)-stained sections. Yellow boxes indicate regions shown at higher magnification. Scale bars, 1 mm (black) and 100 μm (yellow). (b) Principal component analysis (PCA) of RNA sequencing profiles from SMGs of male and female wild-type and *mPR*δ^□^*^/^*^□^ mice (*n* = 3 per group), revealing genotype- and sex-dependent separation. PC1 scores were analyzed using two-way ANOVA followed by simple main-effect analyses. (c) UMAP visualization of epithelial striated duct/granular convoluted tubule (SD/GCT) subclusters from male mice (left). Bar plots showing the proportion of each SD/GCT subpopulation in wild-type and *mPR*δ^□^*^/^*^□^ male mice (right). (d) Total salivary protein concentration in 7-week-old male wild-type and *mPR*δ^□^*^/^*^□^ mice (*n* = 8). (e, f) Proteomic analysis of SMGs from wild-type and *mPR*δ^□^*^/^*^□^ male mice (*n* = 4). (e) Gene Ontology (GO) enrichment analysis of differentially abundant proteins. (f) Heatmap of proteins associated with enriched GO terms. (g, h) Proteomic analysis of saliva from wild-type and *mPR*δ^□^*^/^*^□^ male mice (*n* = 4). (g) GO enrichment analysis of differentially abundant salivary proteins. (h) Heatmap of proteins associated with selected enriched GO terms. (i) Salivary levels of kallikrein 1 (Klk1), TNFα and IgA in wild-type and *mPR*δ^□^*^/^*^□^ male mice (*n* = 8). (j) Plasma TNFα concentrations in wild-type and *mPR*δ^□^*^/^*^□^ male mice (*n* = 8). (k) Representative buccal views of molars and alveolar bone. The distance between the alveolar bone crest (ABC) and cement–enamel junction (CEJ) was measured at the indicated sites (*n* = 7, 9). Scale bar, 1 mm. Data are presented as mean ± SEM. Statistical significance was determined using a two-tailed Student’s *t*-test or one-way ANOVA followed by the appropriate post hoc tests, as indicated (\**P* < 0.05, \*\**P* < 0.01; NS, not significant).

GCT and SD cells constitute the major secretory compartments of the submandibular gland, producing growth factors, antimicrobial proteins and cytokines that are released into saliva^11,15^. Therefore, we next assessed salivary gland function. Although submandibular gland weight, saliva volume, and electrolyte composition were unchanged (Extended Data Fig. 7a–c), total salivary protein content was significantly reduced in mPRδ-deficient males (Fig. 3d). Proteomic analysis of submandibular glands revealed broad reductions in proteins associated with GCT function, secretory vesicles and secretory granules (Fig. 3e, f), indicating impaired secretory capacity. Consistent with this, salivary proteomics demonstrated a marked reduction in proteins involved in proteolysis, wound healing, and responses to external stimuli (Fig. 3g, h), suggesting compromised oral defense. At the molecular level, salivary kallikrein 1 (Klk1) was significantly reduced, whereas the inflammatory markers TNFα and IgA^31^ were increased (Fig. 3i). Plasma TNFα concentrations were likewise elevated (Fig. 3j), indicating that impaired salivary gland function is associated with systemic inflammation.

We challenged mice with models of periodontal disease and metabolic dysfunction to determine whether these defects increase disease susceptibility. mPRδ-deficient males developed significantly more severe periodontal disease than wild-type controls (Fig. 3k). Likewise, mPRδ-deficient males exhibited greater body weight gain and hyperglycemia under high-fat diet conditions, whereas females showed no significant metabolic abnormalities (Extended Data Fig. 7d, e). Collectively, these findings demonstrate that mPRδ is required to maintain salivary gland function and oral homeostasis in a sex-dependent manner, and that its loss predisposes males to oral inflammation and systemic metabolic dysfunction.

### Pregnancy-dependent progesterone–mPR**δ** signaling remodels maternal salivary gland function

mPRδ deficiency produced only modest phenotypes in adult females, likely owing to the limited GCT development. Therefore, we investigated whether pregnancy reactivates progesterone–mPRδ signaling in the maternal submandibular gland. Consistent with this hypothesis, expression of *mPR*δ and epithelial progenitor- and differentiation-associated genes was transiently induced during pregnancy (Fig. 4a). Transcriptomic analysis further revealed that maternal submandibular glands acquired a distinct transcriptional state during gestation, with samples from gestational day 17 (GD17) clearly separated from those of males, non-pregnant females, and postpartum females using PCA (Fig. 4b). This pregnancy-associated transcriptional program was characterized by coordinated induction of GCT-, SD-, and basal-cell-associated genes, consistent with enhanced secretory differentiation (Fig. 4c).

**Fig. 4.**
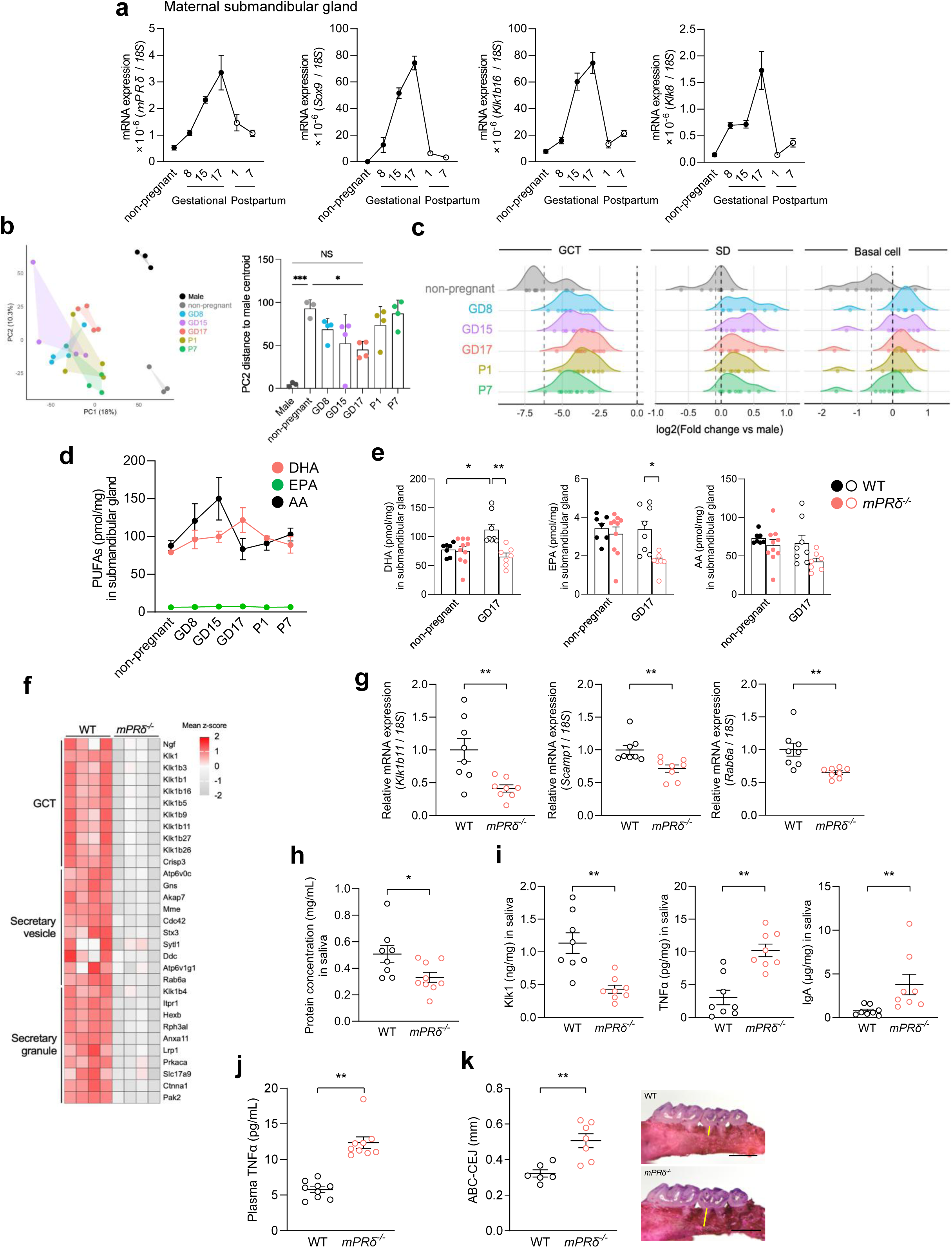
| Pregnancy-dependent progesterone–mPRδ signaling remodels maternal submandibular gland function. (a) mRNA expression of *mPR*δ and epithelial progenitor- and differentiation-associated genes in maternal SMGs across gestation and the postpartum period (*n* = 4). (b) PCA of RNA sequencing profiles from SMGs of male, non-pregnant female, pregnant, and postpartum wild-type mice (left; *n* = 3–4 per group). The distance of each sample from the male centroid along PC2 is shown on the right. GD, gestational day; P, postpartum day. (c) Ridge plots showing the distributions of log□ fold changes relative to male mice for genes associated with GCT, SD, and basal-cell identities in SMGs from non-pregnant, pregnant, and postpartum wild-type mice (*n* = 3–4). Grey dashed lines indicate the mean value for non-pregnant samples. (d) Levels of free DHA, EPA, and AA in wild-type SMGs across the gestational and postpartum periods (*n* = 4). (e) Levels of free DHA, EPA, and AA in SMGs from non-pregnant and GD17 wild-type and *mPR*δ^□^*^/^*^□^ mice (*n* = 7–10). (f) Heatmap of differentially expressed genes associated with GCT function, secretory vesicles, and secretory granules in GD17 SMGs from wild-type and *mPR*δ^□^*^/^*^□^ mice (*n* = 4). (g) RT-qPCR analysis of epithelial differentiation- and secretory-associated genes in GD17 SMGs from wild-type and *mPR*δ^□^*^/^*^□^ mice (*n* = 8). Expression was normalized to 18S rRNA. (h) Total salivary protein concentration in GD17 wild-type and *mPR*δ^□^*^/^*^□^ mice (*n* = 8–9). (i) Salivary Klk1, TNFα, and IgA in GD17 wild-type and *mPR*δ^□^*^/^*^□^ mice (*n* = 8). (j) Plasma TNFα concentration in GD17 wild-type and *mPR*δ^□^*^/^*^□^ mice (*n* = 9). (k) Representative buccal views of molars and alveolar bone. The ABC–CEJ distance was measured at the indicated sites (*n* = 6, 7). Scale bar, 1 mm. Data are presented as mean ± SEM. Statistical significance was determined using a two-tailed Mann–Whitney U test or one-way ANOVA (\**P* < 0.05, \*\**P* < 0.01; NS, not significant).

Given that DHA is the principal downstream effector of progesterone–mPRδ signaling, we examined lipid remodeling during pregnancy. Free DHA levels increased transiently during gestation following the pregnancy-associated increase in AA^32^ (Fig. 4d). This gestational increase in free DHA was significantly attenuated in GD17 mPRδ-deficient mice (Fig. 4e). Consistent with impaired DHA mobilization, RNA sequencing, GO analysis and reverse transcription–quantitative PCR (RT-qPCR) demonstrated coordinated reductions in genes associated with GCT function, secretory vesicles, and secretory granules in GD17 mPRδ-deficient submandibular glands (Fig. 4f, g and Extended Data Fig. 7f), indicating impaired secretory differentiation during pregnancy.

Subsequently, we assessed whether these molecular changes affected maternal salivary gland function. Total salivary protein content was significantly reduced in GD17 mPRδ-deficient mice (Fig. 4h). Likewise, salivary Klk1 levels were reduced, whereas TNFα and IgA levels were increased (Fig. 4i), indicating impaired oral homeostasis. Plasma TNFα concentrations were also elevated (Fig. 4j), demonstrating the presence of systemic inflammation during pregnancy. Consistent with these findings, mPRδ-deficient pregnant mice developed more severe periodontal disease than wild-type controls (Fig. 4k). Furthermore, following maternal periodontal disease induction, offspring born to mPRδ-deficient mothers exhibited reduced body weight after birth (Extended Data Fig. 7g). Collectively, these findings demonstrate that pregnancy transiently activates a progesterone–mPRδ-dependent program that promotes salivary gland secretory differentiation and oral homeostasis. Furthermore, disruption of this pathway predisposes mothers to oral inflammation and adversely affects offspring growth.

### Progesterone–mPR**δ** signaling maintains oral microbial homeostasis and prevents systemic pathology

To determine whether impaired salivary gland function disrupts oral microbial homeostasis, we first tested whether the disease phenotypes observed in mPRδ-deficient mice depended on commensal microorganisms. Antibiotic treatment abolished the differences in periodontal disease severity and metabolic dysfunction between wild-type and mPRδ-deficient mice (Fig. 5a and Extended Data Fig. 8a). Likewise, antibiotic treatment eliminated the elevation of plasma TNFα observed in both adult male and pregnant (GD17) mPRδ-deficient mice (Fig. 5a). These findings demonstrate that the pathological consequences of mPRδ deficiency are microbiota-dependent.

**Fig. 5.**
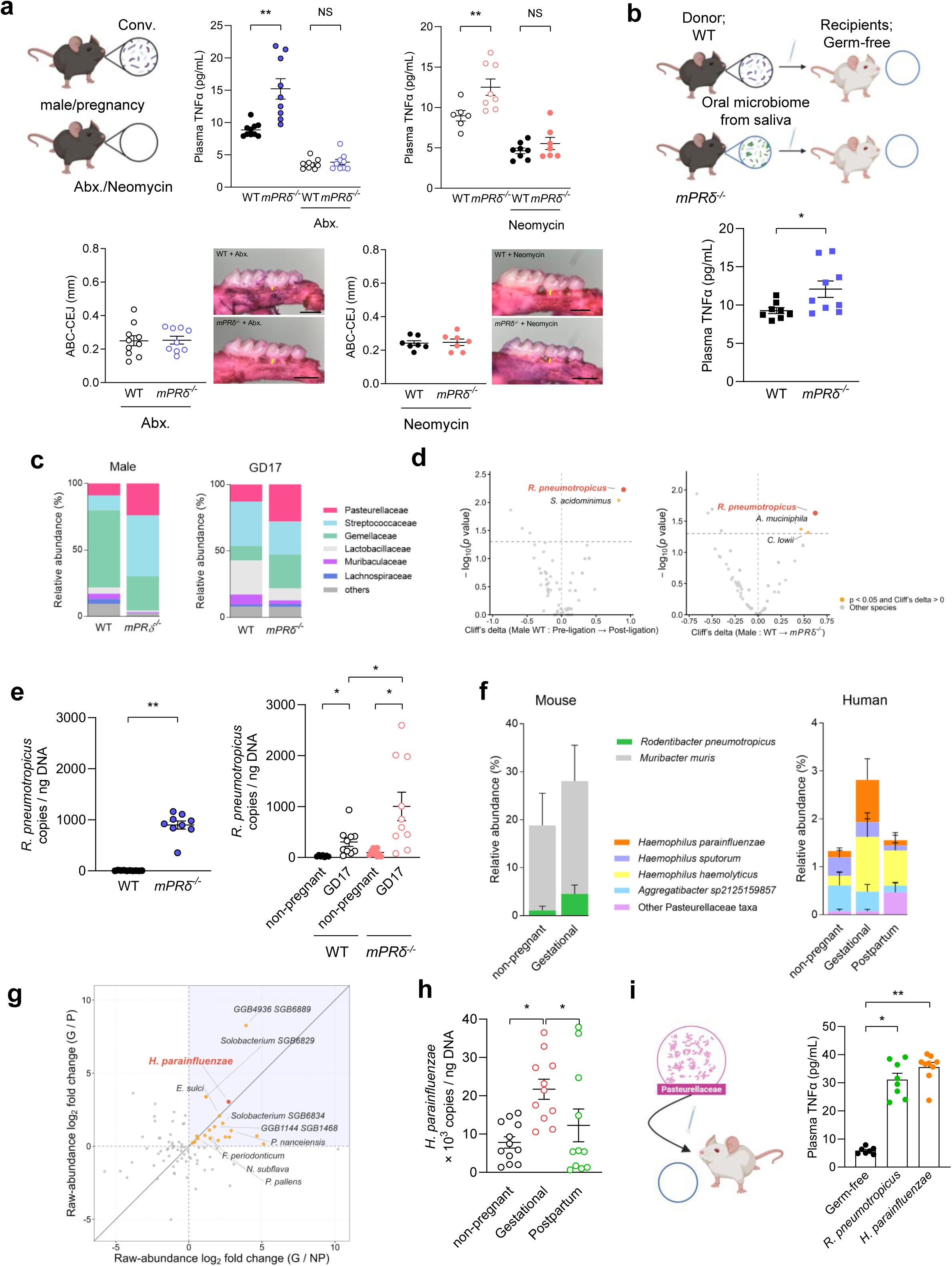
| Progesterone–mPRδ signaling restrains Pasteurellaceae-driven oral dysbiosis and systemic inflammation. (a) Experimental scheme for microbiota depletion (upper left). Plasma TNFα concentrations (upper middle) and periodontal bone loss (lower middle) in wild-type and *mPR*δ^□^*^/^*^□^ adult male mice with or without broad-spectrum antibiotic treatment (*n* = 9–10). Plasma TNFα concentrations (upper right) and periodontal bone loss (lower right) in wild-type and *mPR*δ^□^*^/^*^□^ GD17 pregnant mice with or without neomycin treatment (*n* = 6–8). ABC–CEJ distance and representative buccal views of molars and alveolar bone are shown in the lower panels. Scale bar, 1 mm. (b) Experimental design for transferring oral microbiota from wild-type or *mPR*δ^□^*^/^*^□^ donor mice into germ-free recipients, followed by high-fat-diet feeding for 5 weeks (upper). Plasma TNFα concentrations in recipient mice are shown in the lower panel (*n* = 8–9). (c) Relative abundances of major bacterial families in the oral microbiome of 7-week-old male mice (left) and GD17 pregnant mice (right) in wild-type and *mPR*δ^□^*^/^*^□^ mice (*n* = 4–8). (d) Volcano plots showing bacterial species associated with ligature-induced periodontitis in wild-type adult male mice (left; *n* = 10) and differentially abundant bacterial species between wild-type and *mPR*δ^□^*^/^*^□^ adult male mice (right; *n* = 9–10). The x-axis indicates effect size (Cliff’s delta), and the y-axis indicates −log□(*P* value). (e) Abundance of *Rodentibacter pneumotropicus* in wild-type and *mPR*δ^□^*^/^*^□^ adult male mice (left), and non-pregnant and GD17 pregnant mice (right), quantified by qPCR (*n* = 7–10). (f) Species-level composition of Pasteurellaceae in mouse saliva during non-pregnancy and pregnancy (GD10) (left; *n* = 7–8) and in human saliva during non-pregnancy, pregnancy and the postpartum period (right; *n* = 11–12). (g) Scatter plot comparing pregnancy-associated changes in the abundance of bacterial species in human saliva. The x- and y-axes indicate log□□fold changes during pregnancy relative to the postpartum period (G/P) and non-pregnancy (G/NP), respectively (*n* = 11–12). (h) Abundance of *Haemophilus parainfluenzae* in human saliva during non-pregnancy, pregnancy and the postpartum period, quantified by qPCR (*n* = 11–12). (i) Experimental design for mono-association of germ-free mice with *R. pneumotropicus* or *H. parainfluenzae* (left). Plasma TNFα concentrations in germ-free and mono-colonized mice are shown on the right (*n* = 7–9). Data are presented as mean ± SEM. Statistical significance was determined using a two-tailed Mann–Whitney U-test, except for paired human gestational and postpartum samples, which were analyzed using a two-tailed Wilcoxon signed-rank test (\**P* < 0.05, \*\**P* < 0.01; NS, not significant).

We transplanted oral microbiota from conventionalized donor mice into germ-free (GF) recipients (Extended Data Fig. 8b) to determine whether the altered oral microbiota is sufficient to transmit these phenotypes. Recipients colonized with microbiota from mPRδ-deficient donors exhibited significantly higher plasma TNFα concentrations than those receiving microbiota from wild-type donors (Fig. 5b). Furthermore, microbiota from mPRδ-deficient mice exacerbated high-fat diet-induced metabolic dysfunction in recipient mice (Extended Data Fig. 8c). These findings demonstrate that the altered oral microbiota is sufficient to drive systemic inflammatory and metabolic dysfunction.

We next sought to identify the bacterial taxa responsible for these phenotypes. 16S rRNA gene sequencing revealed marked remodeling of the oral microbiota in mPRδ-deficient mice (Fig. 5c). Members of the family Pasteurellaceae were consistently enriched among the altered taxa, with the greatest expansion observed in both adult male and pregnant mPRδ-deficient mice (Fig. 5c). To identify the bacterial species underlying these changes, we performed long-read sequencing of the mouse oral microbiota. Among all detected oral bacterial species, *Rodentibacter pneumotropicus* was uniquely and consistently enriched following both periodontitis induction and mPRδ deficiency in adult males (Fig. 5d). This species belongs to the family Pasteurellaceae, consistent with the family-level enrichment identified by 16S rRNA sequencing. qPCR further confirmed that *R. pneumotropicus* abundance was significantly increased in both adult males and pregnant mPRδ-deficient mice and was also elevated during pregnancy (Fig. 5e).

To assess the relevance of these findings in humans, we analyzed salivary microbiota from healthy non-pregnant women and a longitudinal pregnancy cohort. Consistent with the mouse data, the abundance of Pasteurellaceae increased significantly during pregnancy. It returned to baseline postpartum (Extended Data Fig. 8d), indicating evolutionarily conserved remodeling of this bacterial family. However, *R. pneumotropicus* was not consistently detected in human saliva. Instead, shotgun metagenomic sequencing identified *Haemophilus parainfluenzae* as the predominant Pasteurellaceae species. It exhibited one of the strongest pregnancy-associated increases among all human oral bacterial species, remaining significantly enriched compared with both non-pregnant and postpartum samples (Fig. 5f–h). Collectively, these findings identify *H. parainfluenzae* as the most likely human functional counterpart of murine *R. pneumotropicus*.

Finally, we mono-colonized GF mice with either *R. pneumotropicus* or *H. parainfluenzae* (Extended Data Fig. 8e) to determine whether these bacteria are sufficient to induce disease phenotypes. Colonization with either species significantly increased plasma TNFα concentrations (Fig. 5i) and exacerbated high-fat diet-induced metabolic dysfunction (Extended Data Fig. 8f). Collectively, these findings establish progesterone–mPRδ signaling as a key regulator of oral microbial homeostasis by restricting the expansion of potentially pathogenic Pasteurellaceae, thereby protecting against oral inflammation and systemic metabolic dysfunction.

### Maternal DHA supplementation rescues mPRδ deficiency–induced defects in oral homeostasis

Finally, we investigated whether DHA, the principal downstream effector of progesterone–mPRδ signaling, could rescue the developmental and functional defects caused by mPRδ deficiency. Pregnant mice were fed a DHA-supplemented diet (Fig. 6a and Supplementary Table 1), which significantly increased free DHA concentrations in maternal plasma and in both maternal and embryonic submandibular glands (Fig. 6b and Extended Data Fig. 8g). DHA supplementation restored pregnancy-associated expression of epithelial differentiation markers, including *Klk1b16*, *Krt14*, and *Klk8*, in the maternal submandibular glands of mPRδ-deficient mice (Fig. 6c).

**Fig. 6.**
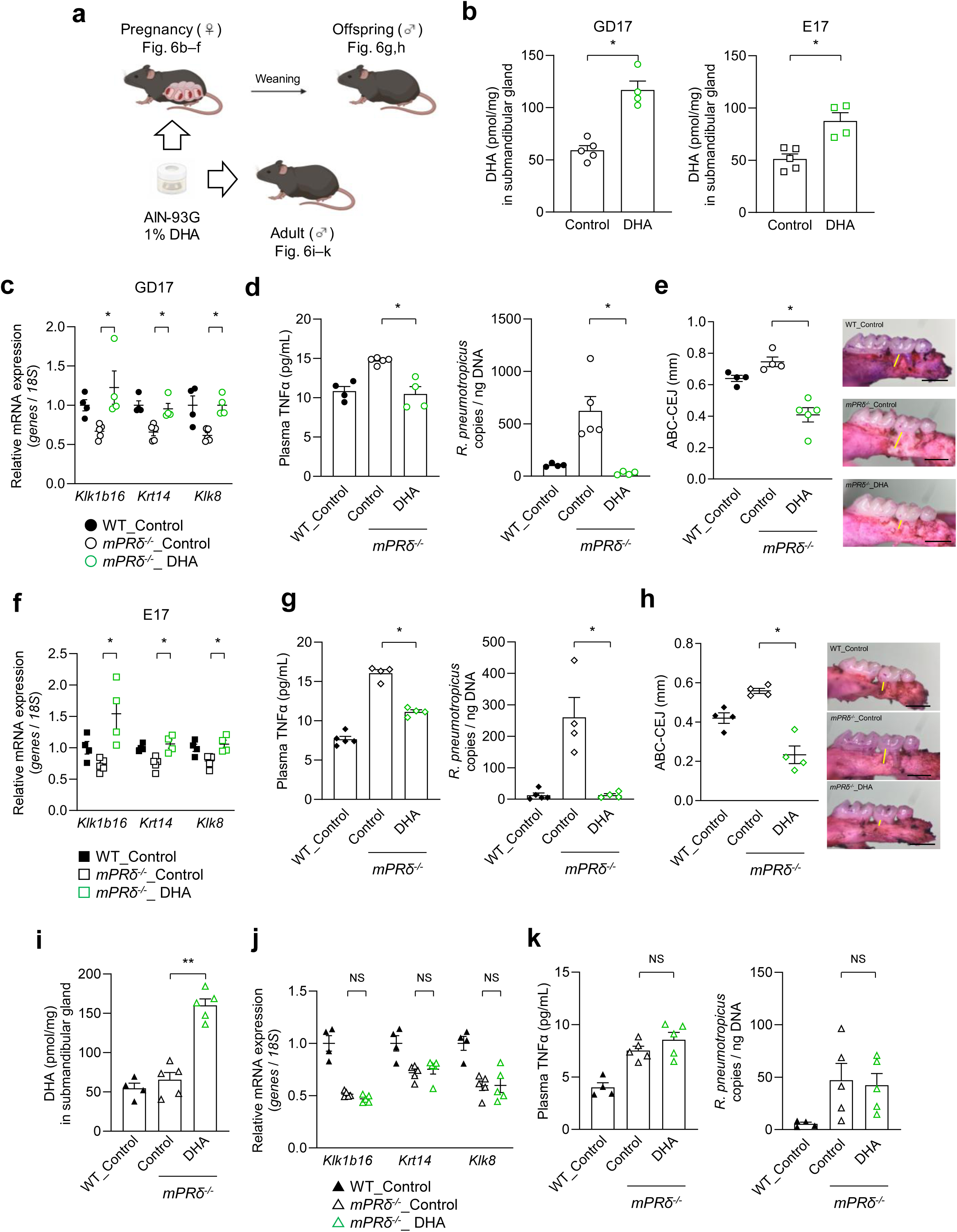
| Maternal DHA supplementation rescues mPRδ deficiency–induced defects in oral homeostasis within a developmental window. (a) Experimental design for DHA supplementation. (b) Free DHA levels in maternal GD17 SMGs and embryonic E17 SMGs from *mPR*δ^□^*^/^*^□^ mice following maternal dietary DHA supplementation (*n* = 4–5). (c) RT-qPCR analysis of *Klk1b16*, *Krt14*, and *Klk8* in maternal GD17 SMGs from wild-type and *mPR*δ^□/□^ mice with or without DHA supplementation (*n* = 4–5). Expression was normalized to 18S rRNA. (d) Plasma TNFα concentrations and oral abundance of *R. pneumotropicus* in GD17 wild-type and *mPR*δ^□/□^ mice with or without maternal DHA supplementation (*n* = 4–5). (e) ABC–CEJ distances and representative buccal views of molars and alveolar bone in GD17 wild-type and *mPR*δ^□/□^ mice with or without maternal DHA supplementation (*n* = 4–5). Scale bar, 1 mm. (f) RT-qPCR analysis of *Klk1b16*, *Krt14*, and *Klk8* in embryonic E17 SMGs from wild-type and *mPR*δ^□/□^ mice with or without DHA supplementation (*n* = 4–5). Expression was normalized to 18S rRNA. (g) Plasma TNFα levels and oral *R. pneumotropicus* abundance in adult male offspring born to control-fed wild-type mothers or to control- or DHA-fed *mPR*δ^□/□^ mothers (*n* = 4–5). (h) ABC–CEJ distances in adult male offspring born to control-fed wild-type mothers or to control- or DHA-fed *mPR*δ^□/□^ mothers (*n* = 4). Scale bar, 1 mm. (i) SMG free DHA levels following direct DHA supplementation in adult male wild-type and *mPR*δ^□/□^ mice (*n* = 4–5). (j) RT-qPCR analysis of *Klk1b16*, *Krt14*, and *Klk8* in SMGs from adult male wild-type and *mPR*δ^□/□^ mice with or without direct DHA supplementation (*n* = 4–5). Expression was normalized to 18S rRNA. (k) Plasma TNFα levels and oral *R. pneumotropicus* abundance in adult wild-type and *mPR*δ^□/□^ male mice with or without direct DHA supplementation (*n* = 4–5). Data are presented as mean ± SEM. Statistical significance was determined using a two-tailed Mann–Whitney U test (\**P* < 0.05, \*\**P* < 0.01).

Consistent with this restoration of glandular differentiation, maternal DHA supplementation normalized plasma TNFα concentrations, suppressed expansion of *R. pneumotropicus*, and significantly ameliorated periodontal disease in pregnant mPRδ-deficient mice, indicating recovery of maternal oral homeostasis (Fig. 6d, e).

Because salivary gland development is initiated during fetal life, we next examined whether maternal DHA supplementation also restored developmental programs in the embryonic submandibular gland. Pregnancy-associated expression of epithelial differentiation markers was rescued in embryonic mPRδ-deficient submandibular glands (Fig. 6f). Strikingly, these developmental effects were accompanied by long-term protection in adult offspring. Maternal DHA supplementation rescued disease phenotypes in adult male offspring, normalizing plasma TNFα concentrations, suppressing *R. pneumotropicus* expansion, and ameliorating periodontal disease (Fig. 6g, h). In contrast, despite markedly increasing free DHA concentrations in plasma and submandibular glands (Fig. 6i and Extended Data Fig. 8h), direct DHA supplementation in adult mPRδ-deficient males (Fig. 6a) failed to restore epithelial differentiation or improve microbial and inflammatory abnormalities (Fig. 6j, k).

Collectively, these findings demonstrate that DHA functions during a critical developmental window to mediate progesterone–mPRδ-dependent programming of salivary gland development and lifelong oral homeostasis. The ability of maternal—but not adult—DHA supplementation to rescue disease establishes developmental programming as the basis for long-term oral microbial homeostasis and systemic health.

## Discussion

Pregnancy is accompanied by profound endocrine remodeling, yet the mechanisms by which maternal hormonal signals shape peripheral organ development and long-term systemic physiology have remained largely unknown. Here, we identify a previously unrecognized progesterone–mPRδ–DHA axis that links pregnancy-associated hormonal changes to salivary gland development, oral microbial homeostasis, and systemic metabolic health. Mechanistically, progesterone–mPRδ signaling selectively mobilizes DHA to promote epithelial differentiation, thereby establishing a developmental program that governs salivary gland function across the life course. Unexpectedly, we further show that this developmental program is transiently reactivated during pregnancy, when it protects maternal oral homeostasis, and that disruption of this pathway remodels the oral microbiota, promotes expansion of potentially pathogenic Pasteurellaceae, and predisposes both mothers and offspring to inflammatory and metabolic dysfunction. Collectively, these findings establish a hormone–lipid–microbiome axis through which maternal endocrine signals program long-term oral and systemic homeostasis.

Mechanistically, our findings identify DHA as the principal downstream effector of progesterone–mPRδ signaling and reveal an unexpected mechanism by which a membrane progesterone receptor selectively mobilizes a specific polyunsaturated fatty acid to regulate epithelial differentiation. Previous studies have implicated mPR signaling in PLA2-dependent lipid remodeling and fatty acid mobilization^24,25,33^. Our findings extend this framework by revealing a striking selectivity of mPRδ signaling for the mobilization of free DHA, consistent with its mobilization from polyunsaturated membrane phospholipid pools. Although modest alterations in several other free fatty acids were detected *in vivo*, these changes were limited in magnitude and are unlikely to account for the broad developmental and physiological phenotypes observed here. Likewise, the substantially lower abundance of DHA-derived lipid mediators relative to DHA itself suggests that DHA, rather than its downstream metabolites, is the principal differentiation-promoting signal. This effect may be mediated through nuclear receptors, such as PPARγ or RXR^34^, or membrane fatty acid receptors, including GPR120 and GPR40^35,36^; however, the precise downstream signaling mechanisms remain to be determined. Importantly, maternal—but not adult—DHA supplementation rescued the epithelial, microbial and inflammatory abnormalities caused by mPRδ deficiency. These findings indicate that DHA acts primarily during a critical developmental window rather than through direct modulation of the oral environment, such as by altering microbiota-derived metabolites or exerting local anti-inflammatory effects^37,38^. Collectively, these findings identify selective DHA mobilization as the central molecular mechanism linking progesterone–mPRδ signaling to developmental programming of salivary gland function and long-term oral homeostasis.

The translational relevance of our findings should be considered in the context of differences in salivary gland organization and sexual dimorphism among species. Although the GCT, a major site of protein secretion in rodents, is absent in humans, analogous secretory functions are thought to be performed by SD compartments^10–15^. Importantly, our data indicated that mPRδ regulates epithelial differentiation at an early stage of ductal lineage specification rather than GCT differentiation per se. This suggests that the underlying developmental mechanism is likely to be conserved despite anatomical divergence between species.

Our findings suggest that pregnancy transiently re-engages a developmental program that is normally active during organogenesis, thereby adapting salivary gland function to the increased physiological demands of gestation. Pronounced sexual dimorphism of the submandibular gland in mice results in highly developed GCT structures in males, whereas females exhibit a transient pregnancy-associated expansion of secretory differentiation. These observations suggest that progesterone–mPRδ signaling is not simply required for gland development but functions as an adaptive mechanism that enhances secretory capacity when physiological demand increases, thereby protecting oral homeostasis during pregnancy. Consistent with this interpretation, alterations in non-secretory SD populations of non-pregnant mPRδ-deficient females suggest that subtle defects may remain latent until challenged by pregnancy.

Finally, the identification of members of the Pasteurellaceae family as the predominant mPRδ-responsive taxa in both mice (*R. pneumotropicus*) and humans (*H. parainfluenzae*) further supports the evolutionary conservation of this host–microbiome interaction. Although the specific bacterial species differ, their shared taxonomic relationship suggests that progesterone-dependent regulation of oral microbial ecology is conserved across mammals. In contrast, the abundance of *Porphyromonas gingivalis*, a major periodontal pathogen in humans^39^, changed only minimally during pregnancy, indicating that pregnancy-associated dysbiosis may initially be driven by microbial taxa other than established periodontal pathogens. Collectively, these findings support the broad translational relevance of the developmental and host–microbiome mechanisms identified in this study.

Our findings further highlight saliva as a key mediator linking developmental programming to oral microbial ecology. Although saliva contains numerous bioactive molecules capable of directly influencing host tissues and microorganisms^11,18^, our results indicated that a major function of saliva in this context is to establish a microbial environment that maintains oral homeostasis. The abolition of the genotype-dependent phenotypic differences by antibiotic treatment, together with their transmission by oral microbiota transfer, demonstrates that microbial communities are both necessary and sufficient to mediate the downstream pathological consequences of mPRδ deficiency.

Rather than causing broad microbial dysbiosis, disruption of progesterone–mPRδ signaling selectively promoted the expansion of members of the Pasteurellaceae family in mice, with a similar pregnancy-associated microbial signature observed in humans. This observation suggests that host developmental programs may regulate specific ecological niches within the oral microbiome rather than globally altering community composition. The ability of *R. pneumotropicus* and *H. parainfluenzae* to recapitulate inflammatory and metabolic abnormalities further identifies these bacteria as functional mediators linking salivary gland dysfunction to systemic disease. More broadly, our findings support a model in which maternal endocrine signals influence long-term systemic physiology by shaping host–microbiome interactions during development.

Our findings also provide a mechanistic framework for understanding the long-recognized association between pregnancy, sexual dimorphism, and oral disease. Although increased susceptibility to gingival inflammation during pregnancy and sex-dependent differences in periodontal disease have been well documented^8–10^, the underlying mechanisms have remained incompletely understood. Our study suggests that progesterone-dependent regulation of salivary gland differentiation shapes the oral microbial environment, thereby influencing susceptibility to inflammatory disease. More broadly, the observation that maternal hormonal signaling programs oral microbial homeostasis and systemic physiology in offspring identifies the salivary gland as a previously unrecognized mediator of developmental programming and intergenerational health.

These findings also have important therapeutic implications. The ability of maternal—but not adult—DHA supplementation to rescue both developmental and adult phenotypes highlights the existence of a critical developmental window during which oral homeostasis can be durably established. Rather than supporting DHA supplementation as a general treatment for established disease, our findings suggest that nutritional or pharmacological interventions targeting the progesterone–mPRδ–DHA pathway during early development may offer new opportunities to prevent disease by targeting developmental programming rather than treating established pathology.

## Limitations of the study

This study has several limitations. First, although our data support a predominant role for the submandibular gland, mPRδ is expressed in multiple tissues, and contributions from other organs cannot be completely excluded. Second, while DHA was identified as the principal downstream effector of mPRδ signaling, the precise intracellular signaling pathways through which DHA promotes epithelial differentiation remain to be defined. Third, although our findings support an evolutionarily conserved host–microbiome interaction, species-specific differences in salivary gland anatomy and oral microbial composition warrant further validation in human systems. Finally, our study focused primarily on oral, inflammatory, and metabolic phenotypes, and whether this developmental pathway contributes to other physiological or pathological processes remains an important question for future investigation.

## Methods

### Animals

Male C57BL/6J mice were purchased from Japan SLC (Shizuoka, Japan; RRID: IMSR_JAX:000664). mPRδ-deficient mice on a C57BL/6J genetic background were generated using embryonic stem cells obtained from the Knockout Mouse Project (KOMP; RRID: MMRRC_059951-UCD). All animals were maintained under standard specific-pathogen-free conditions in a conventional animal facility at 24 °C with a 12-h light/dark cycle. Mice were fed a standard laboratory diet (CLEA Rodent Diet CE-2; CLEA Japan, Inc., Tokyo, Japan) and allowed to acclimate for 1 week prior to experimental procedures. For diet-induced obesity experiments, 7-week-old mice were fed a high-fat diet (HFD) (D12492, 60% kcal fat; Research Diets) for 5 weeks. For DHA supplementation experiments in pregnant mice, pregnant mice were fed an AIN-93G-based diet supplemented with 1% (w/w) DHA (Supplementary Table 1) from GD1 to GD17. The pregnant mice were euthanized on GD17, and E17 fetuses were collected. AIN-93G-fed mothers reared offspring for 4 weeks. After weaning, male mice were fed an AIN-93G diet. For DHA supplementation experiments in adult mice, 7-week-old male mice were fed an AIN-93G-based diet supplemented with 1% (w/w) DHA (Supplementary Table 1) for 14 days. Saliva samples were collected following an established protocol^40^. Briefly, mice were anesthetized and administered pilocarpine hydrochloride (0.75 mg/kg for proteomics and 1 mg/kg for electrolyte analysis; Nacalai Tesque, Kyoto, Japan, #2800831) via intraperitoneal injection to induce salivation. Saliva was absorbed for 30 minutes using cylindrical absorbent swabs and subsequently recovered using centrifugation at 7,500 × g for 2 minutes at 4 °C. The volume of collected saliva was measured, and samples were stored at −80 °C until subsequent analyses. All animal experiments were approved by the Kyoto University Animal Experimentation Committee (approval number: Lif-K26002) and Tokyo University of Agriculture and Technology (permit number: R08-49) and were conducted in accordance with institutional guidelines to minimize animal suffering.

### RNA extraction and RT-qPCR

Total RNA was extracted using the RNeasy Mini Kit (Qiagen, Hilden, Germany), RNAiso Plus (TAKARA, Shiga, Japan), or Sepasol-RNA I Super G (Nacalai Tesque), following the manufacturers’ protocols. Complementary DNA (cDNA) was synthesized from total RNA using Moloney murine leukemia virus reverse transcriptase (Invitrogen, Carlsbad, CA, USA). RT–qPCR was performed using SYBR Premix Ex Taq II (TAKARA) on a StepOnePlus Real-Time PCR System (Applied Biosystems, Foster City, CA, USA). The thermal cycling program consisted of an initial denaturation step at 95 °C for 30 seconds, followed by 40 cycles of denaturation at 95 °C for 5 seconds, annealing at 58 °C for 30 seconds, and extension at 72 °C for 1 minute. Melt curve analysis was conducted with sequential incubations at 95 °C for 15 seconds, 60 °C for 1 minute, and 95 °C for 15 seconds. The primer sequences used were as described in Supplementary Table 2.

### Biochemical analyses

Saliva and plasma protein levels (Pierce BCA Protein Assay Kit; ThermoFisher Scientific, Waltham, MA, USA), TNFα (Mouse TNF-alpha Quantikine ELISA Kit; R&D Systems, Minneapolis, MN, USA), IgA (Mouse IgA ELISA Kit; Bethyl Laboratories, Inc., Montgomery, TX, USA), non-esterified fatty acids (LabAssay NEFA; Wako Pure Chemical Co. Ltd., Osaka, Japan), triglycerides (LabAssay Triglyceride; Wako Pure Chemical Co. Ltd.), and total cholesterol (LabAssay Cholesterol; Wako Pure Chemical Co. Ltd.) in mice were measured following the manufacturer’s protocols. Salivary electrolytes were measured by Oriental Yeast Co., Ltd (Tokyo, Japan) using an enzymatic assay for Ca²□ and an ion-selective electrode method for the other ions. Blood glucose levels were assessed with a handheld glucose meter (OneTouch Ultra; LifeScan, Milpitas, CA, USA).

### Steroid measurements

Plasma samples were subjected to steroid extraction by mixing with methanol containing an internal standard, followed by the addition of ethyl acetate. After centrifugation at 8,000 × *g* for 15 minutes at 4 °C, the organic phase containing steroids was collected and evaporated to dryness. The resulting residues were reconstituted in methoxyamine hydrochloride (20 mg/mL in pyridine) and incubated at 60 °C for 45 minutes to allow derivatization. After drying, samples were resuspended in acetonitrile and analyzed using liquid chromatography–tandem mass spectrometry (LC–MS/MS). Chromatographic separation was performed on an ACQUITY UPLC BEH C18 column (2.1 × 150 mm, 1.7 μm; Waters Corporation, Milford, MA, USA) using a methanol gradient in 10 mM ammonium formate aqueous solution. LC–MS/MS analysis was carried out on an Acquity UPLC system coupled to a Xevo TQD triple quadrupole mass spectrometer (Waters Corporation) as described previously^25^.

### Primary culture of submandibular gland cells

Timed-pregnant C57BL/6J mice were obtained from Japan SLC, and embryonic day 18 (E18) submandibular glands (SMGs) were isolated. Briefly, SMGs were carefully dissected using sterile forceps, washed with 1× PBS, and finely minced in culture medium. The medium consisted of Advanced Dulbecco’s Modified Eagle Medium (DMEM) (Thermo Fisher Scientific) supplemented with 1% HEPES, 1% GlutaMAX, and 1% penicillin–streptomycin. Tissues were incubated in medium containing collagenase II (0.6 mg/mL) (CLS-2; Worthington Biochemical Corporation, Lakewood, NJ, USA) and hyaluronidase (0.5 mg/mL) (HSE-300; Worthington Biochemical Corporation) with 6.25 mM CaCl□ at 37 °C for 30 minutes to isolate epithelial cells. After mechanical dissociation, the suspension was passed through a 100 μm mesh filter and centrifuged at 300 × *g* for 3 minutes. The cell pellet was then resuspended in TrypLE (Thermo Fisher Scientific) and incubated at 37 °C for 15 minutes. Following further mechanical dissociation, the suspension was filtered through a 40 μm mesh and centrifuged at 300 × *g* for 3 minutes. The resulting cells were washed with culture medium. For intracellular polyunsaturated fatty acid (PUFA) quantification, cells were seeded in 24-well plates at a density of 1.7 × 10□ cells per well on collagen I–coated plates (50 μg/mL; rat tail collagen type I, Sigma Aldrich). Cells were cultured in the adjusted medium [advanced DMEM/F-12 medium (Life Technologies, Waltham, MA, USA) supplemented with 1% penicillin–streptomycin, GlutaMAX (Life Technologies), 10 mM HEPES (Nacalai Tesque), 1% (v/v) N2 supplement (Life Technologies), 1% (v/v) B27 supplement (Life Technologies), Wnt3a-conditioned medium (Wnt3a-CM), dexamethasone (1 μM), Y-27632 (10 μM), and growth factors including EGF (50 ng mL□¹; PeproTech, Cranbury, NJ, USA), FGF (10 ng mL□¹; FGF max, Sigma Aldrich), IGF-1 (100 ng mL□¹), insulin (10 μg mL□¹), and R-spondin (1 μg mL□¹; Sigma Aldrich)]. Cells were treated with progesterone for 18 hours, followed by methanol fixation. For organoid culture, epithelial cells were isolated from embryonic SMGs using fluorescence-activated cell sorting (FACS) (Extended Data Fig. 3a). Single-cell suspensions were stained with the following antibodies for 15 minutes at 4 °C: PE-Cy7-conjugated anti-CD31 (BioLegend, San Diego, CA, USA), PE-Cy7-conjugated anti-CD45 (BioLegend), and APC-conjugated anti-EpCAM (BioLegend). Cells were washed with sorting buffer and sorted using FACSMelody and FACSChorus (BD Biosciences, Franklin Lakes, NJ, USA). Cells were first gated based on FSC and SSC parameters to exclude debris, followed by exclusion of dead cells using 7-AAD staining. Target epithelial cells were isolated as EpCAM-positive and CD31/CD45-negative cells. The purity of sorted cells exceeded 95%, and data were analyzed using FlowJo v10 (BD Biosciences). Isolated epithelial cells were centrifuged at 500 × *g* for 5 minutes at 4 °C, mixed with Matrigel, and plated at 50,000 cells per well in a 24-well plate in culture medium. Organoids were maintained at 37□°C in 5% CO□ as described previously^41^. Organoids were continuously treated with progesterone (100 nM) or PUFAs (10 μM each) for 9 days and were passaged on day 6 while maintaining the respective treatments. Total RNA was extracted using the RNeasy Mini Kit (Qiagen).

### Histological analysis

SMGs were excised and fixed overnight at 4□°C in 10% neutral buffered formalin. Fixed tissues were processed, embedded in paraffin, and sectioned at 5□μm thickness using a microtome. Sections were stained with hematoxylin and eosin (H&E) using standard procedures.

### *In situ* hybridization

Mouse embryos were rapidly frozen in powdered dry ice, and cryosections of 16 μm thickness were prepared using a cryostat. Sections were stored at −80 °C until use. A ^35^S-labeled antisense RNA probe specific for mouse mPRδ was synthesized using *in vitro* transcription with T7 RNA polymerase in the presence of uridine 5′-α-[^35^S] thiotriphosphate (GE Healthcare, Chicago, IL, USA). *In situ* hybridization was performed on tissue sections using the radiolabeled probe as previously described^27^. Following hybridization, sections were exposed to X-ray film (BioMax MR; Kodak, Rochester, NY, USA) for 10 days. After autoradiographic exposure, sections were counterstained with H&E for histological visualization.

### RNA sequencing

Total RNA was isolated from submandibular organoids or SMGs of mice using RNAiso Plus reagent (TAKARA) in combination with the RNeasy Mini Kit (Qiagen). RNA integrity, quality, and concentration were assessed with an Agilent 2100 Bioanalyzer using the RNA 6000 Nano Kit (Agilent, Santa Clara, CA, USA). Strand-specific RNA sequencing libraries were generated using the NEBNext Ultra II Directional RNA Library Prep Kit for Illumina together with NEBNext Multiplex Oligos for Illumina (Dual Index Primers Set 1). Libraries were sequenced on an Illumina NovaSeq 6000 platform to produce paired-end reads of 150 bp, yielding approximately 4 Gb of data per sample.

Raw sequencing reads were processed with Trimmomatic (v0.39) to remove adaptor sequences and low-quality bases. The quality of the filtered reads was evaluated using FastQC (v0.11.8-2). Cleaned reads were aligned to the mouse reference genome (NCBI GRCm39) using STAR (v2.7.10a). Gene-level expression counts were normalized by relative log expression, and differential expression analyses were performed using RSEM (v1.3.3) and edgeR. Differentially expressed genes (DEGs) were defined as those with a Benjamini–Hochberg false discovery rate (FDR)–adjusted *P* < 0.1. Subsequent GO enrichment analysis of these DEGs defined statistical significance at an adjusted *P* < 0.05, as described previously^25^. PCA plots were generated using the top 10,000 most variable genes. The marker genes analyzed in the ridge plots were identical to those utilized in Figure 1f (comprising 10–11 genes per group).

### Single-cell RNA sequencing

SMGs were collected from three P49 mice and ten P0 mice. Single-cell suspensions were generated from fresh-frozen SMG tissues following fixation using the Chromium Next GEM Single Cell Fixed RNA Preparation Kit (10x Genomics, Pleasanton, CA, USA), in accordance with the manufacturer’s user guide (CG000553). Single-cell RNA sequencing libraries were prepared using the Chromium Next GEM Single Cell Fixed RNA Sample Preparation Kit (CG000527, Rev. E; 10x Genomics). Libraries were sequenced on a NovaSeq X Plus system (Illumina, San Diego, CA, USA). Sequencing data were processed using Cell Ranger (v7.1.0) for read alignment, barcode assignment, and unique molecular identifier (UMI) counting. The resulting gene–cell count matrices were imported into R (v4.5.1) for downstream analysis using the Seurat package (v5.4.0). Quality control filtering was applied to exclude low-quality cells, including those with mitochondrial gene-derived UMIs exceeding 5%, as well as cells with fewer than 300 detected genes or more than 4,500 genes, thereby removing non-cellular events and potential doublets prior to further analyses.

### Proteome analysis

For proteomic analysis, submandibular gland tissues were lysed using the EasyPep Magnetic MS Sample Prep Kit according to the manufacturer’s instructions. Protein concentration was determined using a bicinchoninic acid (BCA) assay. For proteomic analysis, 50 µg of protein per sample was processed for enzymatic digestion and peptide cleanup using the same kit, and 200 ng of peptides were subjected to LC–MS/MS analysis. Saliva samples were subjected to SDS–PAGE under reducing conditions, followed by silver staining. Gel bands excluding the region around 45 kDa (amylase) were excised, destained, and digested with trypsin in Tris buffer (pH 8.0) at 37 °C for 20 hours. Peptides were extracted, desalted, and quantified, and 500 ng of peptides were subjected to LC–MS/MS analysis. Peptides were desalted using EASY-Spray PepMap cartridges and analyzed by LC–MS/MS using a Vanquish Neo coupled to an Orbitrap Exploris 480 (Thermo Fisher Scientific) in data-independent acquisition (DIA) mode, with nanoflow-LC electrospray ionization in positive mode. Raw data were processed using DIA-NN (v1.9) and searched against the UniProtKB/Swiss-Prot database with standard parameters. Precursor identifications were filtered at FDR of <1%. Contaminant proteins were excluded from the analysis, and missing values were imputed using a minimum value approach. Submandibular gland samples were normalized using the median method, and proteins with an adjusted *P* < 0.1 were considered significant. Saliva samples were normalized using quantile normalization, and proteins with *P* < 0.05 were considered significant. Differential expression analysis was performed using the limma package in R, followed by GO enrichment analysis. Heatmaps were generated using median-normalized values.

### Oral microbial characterization

Samples were collected from the murine oral cavity using sterile ultra-fine swabs for oral microbiome profiling. Genomic DNA was extracted from frozen samples with the QIAamp UCP Pathogen Kit (QIAGEN) following established procedures. Partial 16S rRNA gene fragments encompassing the V3-V4 hypervariable region were amplified using the primer pair 341F (5′-TCGTCGGCAGCGTCAGATGTGTATAAGAGACAGCCTACGGGNGGCWGCAG-3′) and 805R (5′-GTCTCGTGGGCTCGGAGATGTGTATAAGAGACAGGACTACHVGGGTATCTAATCC-3′). PCR amplicons were purified using AMPure XP beads (Beckman Coulter, Brea, CA, USA) and sequenced on an Illumina MiSeq platform with the MiSeq Reagent Kit v3 (600-cycle configuration). Raw paired-end sequencing reads were processed using QIIME 2 (v2026.4). Sequence data were imported using the PairedEndFastqManifestPhred33V2 format, and primer sequences were removed using the q2-cutadapt plugin. Quality filtering, denoising, paired-end read merging, and chimera removal were performed using the q2-dada2 plugin. Amplicon sequence variants (ASVs) generated by DADA2 were used for subsequent analyses. Taxonomic classification was performed using the q2-feature-classifier plugin with a scikit-learn naïve Bayes classifier trained specifically on the 341F–805R region of the SILVA 138.2 SSURef NR99 database.

For long-read sequencing analysis, full-length 16S rRNA genes were amplified using the PacBio barcoded forward (5′-Phos-GCATCNNNNNNNNNNAGRGTTYGATYMTGGCTCAG-3′) and barcoded 1492Rmod (5′-Phos-GCATCNNNNNNNNNNRGHTACCTTGTTACGACTT-3′) primers; ‘Phos’ indicates a 5′-phosphate modification, and ‘N’ represents a unique PacBio barcode sequence for each sample. Subsequent procedures followed the manufacturer’s protocol. The sequencing library was prepared using the PacBio SMRTbell Prep Kit 3.0 according to the manufacturer’s protocol. Sequencing was performed on the PacBio Revio system with the option full resolution base qual = TRUE. PacBio HiFi reads were generated automatically using SMRT Link software (v25.1) with default parameters. Read segmentation was conducted using Skera (v1.3.0), and demultiplexing was performed with Lima (v2.12.0) under the HIFI-ASYMMETRIC preset. Full-length 16S rRNA gene ASVs were inferred from demultiplexed HiFi reads using the DADA2 package^42^ (v1.30.0) in R (v4.3.3) according to the previously described DADA2 for PacBio workflow^43^ with slight modifications. The reads were subjected to quality filtering and trimming using the filterAndTrim function with the following parameters: minQ = 3, minLen = 1300, maxLen = 1600, maxN = 0, rm.phix = FALSE, maxEE = 5. ASVs were then subjected to a homology search against RefSeq16S rRNA sequences (downloaded on 24 June 2025) using BLASTN (v2.16+), with a maximum E-value cut-off of 1 × 10^-10^. The highest bitscore determined top hits. ASVs with alignment coverage of ≥90% were taxonomically assigned and summarized at different taxonomic levels using identity thresholds of 70% for phylum, 94% for genus, and 97% for species.

qPCR was performed using SYBR Premix Ex Taq II (TAKARA) with a StepOnePlus Real-Time PCR System (Applied Biosystems). The bacterial primer sequences used in this study are listed in Supplementary Table 3. *R. pneumotropicus* and *H. parainfluenzae* were used as reference strains for calibration in DNA-based quantification of bacterial numbers.

### Shotgun metagenomic sequencing data analysis

DNA extracted from human saliva was quantified using the Qubit dsDNA HS Assay Kit (Thermo Fisher Scientific). Sequencing libraries were prepared from 100 ng of DNA using the Illumina DNA Prep kit with Illumina DNA/RNA UD Indexes (Illumina), following the manufacturer’s instructions. Briefly, genomic DNA was fragmented and adapter-tagged via on-bead tagmentation, followed by library amplification and purification. Sequencing was performed on the NextSeq 2000 platform (Illumina) using the NextSeq 1000/2000 P2 XLEAP-SBS Reagent Kit (600 cycles), with PhiX Control (Illumina) spiked in as an internal standard. A minimum of 20 million paired-end reads per library was targeted. For bioinformatic analysis, paired-end reads were merged using BBmerge (v39.06) and host-derived sequences were removed using Kneaddata (v0.12.2) against the GRCh37-based hg37_v0.1 human-contamination reference database. Taxonomic profiling was conducted using MetaPhlAn (v4.0.6) with the mpa_vJun23_CHOCOPhlAnSGB_202403 database^44^.

### Human salivary sample collection

Samples were collected between April 2025 and June 2026. All experimental procedures involving human salivary sample collection were performed according to the protocols approved by the Ethics Committee of Kyoto University (approval numbers: R2875-7) and Kumamoto University (approval numbers: 2892) and were conducted in accordance with their guidelines. All participants provided written informed consent. Participants included Japanese non-pregnant and pregnant women aged ≥18 years receiving obstetric care for pregnancy and delivery, and patients undergoing surgery for gynecological diseases. Participants unable to provide informed consent or deemed unsuitable by the investigators were excluded. Salivary samples from non-pregnant and pregnant participants were collected before and after delivery using salivary collection tubes and stored at −80 °C until processing and analysis.

### Bacterial strains and culture

*R. pneumotropicus* JCM14074T was obtained from the Japan Collection of Microorganisms (JCM), RIKEN BRC (Tokyo, Japan), and cultured according to the recommended conditions.. *H. parainfluenzae* was isolated from human saliva by serial dilution and cultivation on Gifu anaerobic medium (GAM) agar supplemented with 5% (v/v) defibrinated horse blood and vancomycin at a final concentration of 5 μg/ml. Plates were incubated at 37□°C under 5–10% CO□ for 24–48□h. Selected colonies were purified by repeated streaking on the same medium, and species identity was confirmed by Sanger sequencing of PCR amplicons targeting the 16S rRNA gene and *rpoB*.

### Non-esterified fatty acid measurements

Samples were supplemented with an internal standard (C17:1) and homogenized in methanol (1 mL), followed by lipid extraction with chloroform (2 mL) and 0.5 M potassium chloride (0.75 mL). After phase separation, the organic lipid fraction was collected, evaporated to dryness, and reconstituted in chloroform:methanol (1:3, v/v) for analysis. Non-esterified fatty acid (FA) profiling was performed using liquid chromatography–tandem mass spectrometry (LC–MS/MS) with an LCMS-8045 triple quadrupole mass spectrometer (Shimadzu, Kyoto, Japan). Chromatographic separation was achieved on a Shim-pack Sceptor C18 column (2.1 × 150 mm, 1.9 μm; Shimadzu) with an acetonitrile gradient in 10 mM ammonium formate aqueous solution at a flow rate of 0.4 mL/min, with the column temperature maintained at 50 °C. Mass spectrometric detection was conducted in negative ionization mode with a capillary voltage of 3,000 V. The desolvation and source temperatures were set to 500 °C and 150 °C, respectively. Multiple reaction monitoring (MRM) transitions were individually optimized for each target FA using authentic standards^25,37^.

### Comprehensive analysis of lipid mediators

Lipid mediators were quantified from tissue samples using solid-phase extraction with Oasis HLB cartridges (1 mg; Waters Corporation), following an established protocol. Tissues were homogenized in methanol, and the resulting supernatants were diluted with water to adjust the methanol concentration to approximately 7%. Diluted extracts were loaded onto preconditioned cartridges and sequentially washed with 0.1% formic acid, 15% ethanol, and hexane. Lipid mediators were eluted with 200 μL of methanol, and eluates were evaporated to dryness. The residues were reconstituted in 20 μL of methanol and subjected to liquid chromatography–tandem mass spectrometry (LC–MS/MS) analysis. A broad spectrum of lipid mediators was analyzed using the LC/MS/MS Method Package for Lipid Mediators ver. 2 (Shimadzu) on an LCMS-8060NX triple quadrupole mass spectrometer (Shimadzu). Chromatographic separation was performed on a Kinetex C8 column (2.1 × 150 mm, 2.6 μm; Phenomenex, Torrance, CA, USA)^25^.

### Phospholipid measurements

Total lipids were extracted from SMG samples using the Bligh and Dyer method. The lipid extracts were dried using a centrifugal evaporator, reconstituted in methanol:isopropanol (1:1, v/v), and stored at −20 °C until analysis. Total phospholipid content was determined by phosphorus assay. Samples were then dried under a stream of nitrogen at room temperature and reconstituted in isopropanol:methanol (1:1, v/v) containing internal standards, including 12:0/12:0 phosphatidylcholine (PC), phosphatidylethanolamine (PE), phosphatidylserine (PS), phosphatidylglycerol (PG), phosphatidic acid (PA), and 8:0/8:0 phosphatidylinositol (PI), to achieve a final phospholipid concentration of 100 μM.

Lipidomic analyses were performed by liquid chromatography–electrospray ionization mass spectrometry (LC–ESI–MS) using a Shimadzu Nexera UPLC system (Shimadzu) coupled to a QTRAP 4500 hybrid triple-quadrupole linear ion-trap mass spectrometer (SCIEX). Chromatographic separation was carried out on a SeQuant ZIC-HILIC PEEK-coated column (250 mm × 2.1 mm, 1.8 μm; Millipore) maintained at 50 °C. Mobile phase A consisted of water:acetonitrile (95:5, v/v) containing 10 mM ammonium acetate, and mobile phase B consisted of water:acetonitrile (50:50, v/v) containing 20 mM ammonium acetate. The gradient program was as follows: 0–22 min, 0–40% B; 22–25 min, 40% B; and 25–30 min, 0% B, at a flow rate of 0.3 mL/min. Instrument parameters were set as follows: curtain gas, 30 psi; collision gas, 7 arbitrary units; ion-spray voltage, −4,500 V; source temperature, 700 °C; ion source gas 1, 30 psi; and ion source gas 2, 70 psi. Phospholipid species were detected by multiple reaction monitoring as previously described^25^. For semi-quantitative comparison across phospholipid classes, the abundance of each phospholipid species was normalized to the total PC content within the same sample and expressed as a percentage of total PC.

### HEK293 cells expressing mouse mPR**δ**

Flp-In T-REx HEK293 cells were obtained from Invitrogen and used to generate a stable cell line expressing mouse mPRδ. Cells were co-transfected with pcDNA5/FRT/TO-E-tag-mPRδ and pOG44 plasmids using Lipofectamine (Invitrogen). Following transfection, cells were selected and maintained in DMEM supplemented with 10% fetal bovine serum (FBS), 10 μg/mL blasticidin S (Funakoshi, Tokyo, Japan), and 100 μg/mL hygromycin B (Gibco, Grand Island, NY, USA) as previously described^22,25^. Cultures were maintained at 37 °C in a humidified incubator with 5% CO and used under the indicated experimental conditions. To quantify the intracellular PUFAs, HEK293 cells were plated in 24-well plates (1.7 × 10^5^ cells/well). Cells were treated with progesterone and, where indicated, doxycycline (10 μg/mL) to induce mPRδ expression simultaneously for 18 hours, after which they were fixed in methanol and stored at −80 °C until analysis. For intracellular calcium ([Ca²□]□) measurements, cells were seeded onto black 96-well plates at a density of 3 × 10□ cells per well and cultured for 24 hours. The cells were incubated for an additional 24 hours with doxycycline (10 μg/mL). Cells were then loaded with calcium-sensitive dye by incubation for 1 hour at 37 °C in Hank’s balanced salt solution (HBSS; pH 7.4) containing component A of the calcium assay kit (Molecular Devices, Sunnyvale, CA, USA). Progesterone solutions prepared in HBSS were dispensed into a separate 96-well plate and applied during real-time fluorescence measurements using a FlexStation 3 Multi-Mode Microplate Reader (Molecular Devices) to monitor [Ca²□]□ mobilization. For cAMP measurements, HEK293 cells were plated in 24-well plates and cultured for 24 hours, followed by induction with doxycycline (10 μg/mL) for an additional 24 hours. Cells were lysed in 0.1 N HCl, and intracellular cAMP levels were quantified by enzyme immunoassay using a cAMP EIA kit (Cayman Chemical, Ann Arbor, MI, USA) according to the manufacturer’s instructions. All experiments were performed in duplicate.

### Ligature-induced periodontal disease model

Experimental periodontal disease was established by ligature placement according to a previously reported method^45^. 7-week-old male and GD10.5 pregnant mice were anesthetized with a mixture of three agents: medetomidine (0.3 mg/kg; Meiji Animal Health, Tokyo, Japan), midazolam (4.0 mg/kg; Sandoz, Tokyo, Japan), and butorphanol (5.0 mg/kg; Meiji Animal Health). A 6-0 silk ligature (F.S.T., Foster City, CA, USA) was carefully tied around the maxillary second molar. After the procedure, anesthesia was reversed by administration of atipamezole (3.0 mg/kg; Nippon Zenyaku Kogyo, Fukushima, Japan). To measure alveolar bone resorption, alveolar bone loss was evaluated 7 days after ligature placement by measuring the distance between the cementoenamel junction (CEJ) and the alveolar bone crest (ABC), based on a previously described method with minor modifications^45^. Briefly, harvested mouse skulls were autoclaved to facilitate removal of soft tissues, which were subsequently cleared manually. The specimens were then treated with 3% H□O□for 1 hour for bleaching and sterilization. Following this step, the skulls were stained with Eosin Y and Phloxine B, as well as 0.05% Toluidine Blue (Muto Pure Chemicals, Tokyo, Japan). Images of the stained samples were captured using a stereomicroscope (Micronet, Saitama, Japan) together with a ruler (KENIS LIMITED, Osaka, Japan) for calibration. The ABC–CEJ distance was quantified using ImageJ software (National Institutes of Health, Bethesda, MD, USA).

### Antibiotic treatment

To deplete the oral microbiota, mice were treated with antibiotics before the indicated experiments. Adult male mice received a broad-spectrum antibiotic cocktail consisting of ampicillin (0.5 g/L, Wako Pure Chemical Co. Ltd.), vancomycin (0.2 g/L, Wako Pure Chemical Co. Ltd.), neomycin sulfate (0.5 g/L, Wako Pure Chemical Co. Ltd.), and metronidazole (0.5 g/L, Wako Pure Chemical Co. Ltd.) in the drinking water for 4 weeks. Pregnant mice received neomycin (1 g/L) in the drinking water from gestational day 0 (GD0) until the indicated experimental endpoint. Antibiotic-containing water was replaced every 2–3 days^27^.

### GF and gnotobiotic experiments

Germ-free (GF) ICR mice were housed in vinyl isolators under a 12-hour light/dark cycle. Sterility was regularly confirmed using both aerobic and anaerobic culture methods, in addition to qPCR targeting the 16S rRNA gene with freshly collected fecal samples^44^. The mice were provided with a chow diet (CL-2, CLEA; irradiated at 50□ kGy) and had free access to autoclaved water. Oral swab samples collected from wild-type or mPRδ-deficient male donor mice were suspended in 200 μL of fresh PBS, and the suspensions were administered once to the oral cavity of 7-week-old GF ICR male recipient mice by swabbing to establish oral microbiota-colonized mice^46^. Swabbing was performed only once. After colonization, each gnotobiotic mouse was housed in an independent vinyl isolator and fed an irradiated HFD (50 kGy) for 5 weeks in diet-induced obesity experiments^44^. In mono-colonized experiments, 7-week-old GF ICR male mice were colonized by swabbing with each bacterial strain (*R. pneumotropicus* or *H. parainfluenzae*; 1 × 10^8^ CFU in 200□μL fresh PBS). After colonization, each mono-colonized mouse was housed in an independent vinyl isolator and fed an HFD (50 kGy irradiated) for 5 weeks in diet-induced obesity experiments^44^.

### Statistical analysis

All quantitative results are presented as mean values with the corresponding standard error of the mean (SEM). Statistical evaluations were carried out using GraphPad Prism software (GraphPad Software Inc., La Jolla, CA, USA; RRID: SCR_002798). The distribution of each dataset was examined for normality using the Shapiro–Wilk test. For comparisons between two groups, either a two-tailed Student’s *t*-test or a two-tailed Mann–Whitney U-test was applied, depending on whether the data satisfied the assumption of normality. For experiments involving multiple factors, two-way analysis of variance (ANOVA) followed by Bonferroni’s multiple-comparison test was performed. For comparisons among three or more groups, one-way ANOVA followed by the appropriate multiple-comparison test was used. Paired human pregnancy and postpartum samples were analyzed using a two-tailed Wilcoxon signed-rank test. Effect sizes for microbiome comparisons were calculated using Cliff’s delta. Multiple-testing correction was performed as described for each omics analysis. *P* < 0.05 was considered statistically significant.

## Data availability

Bulk RNA-seq and single-cell RNA-seq datasets have been deposited in the DDBJ under BioProject accession PRJDB40530 (raw data: DRA026633, DRA026635, DRA026636, DRA029216, DRA029471; processed data: E-GEAD-1264, E-GEAD-1265, E-GEAD-1266, E-GEAD-1292, E-GEAD-1297). Microbiome sequencing datasets have been deposited under BioProject accessions PRJDB42895, PRJDB42896 (raw data: DRA029522, DRA029523, DRA029550, DRA029551). Proteomics data have been deposited in PRIDE under accession PXD080362. All other data relevant to this study have been deposited in the Dryad database (DOI: 10.5061/dryad.vx0k6dk7n).

## Materials availability

All biological materials generated in this study are available from the corresponding author upon reasonable request.

## Acknowledgments

### Funding

This work was supported by research grants from the AMED (JP23gm1510011 to I.K.), JST-MOONSHOT (JPMJMS2023 to I.K. and J.A.), and JSPS KAKENHI (JP24KJ1480 to M.Yamano, JP25H01428 to J.M.).

### Author contributions

M.Yamano performed the experiments, interpreted the data, and wrote the manuscript; J.M. performed the experiments, interpreted the data, and wrote the manuscript; D.S. performed the experiments and wrote the manuscript; Y.M. performed the experiments and interpreted the data; T.I. performed the experiments and interpreted the data; R.O.K. performed the experiments and interpreted the data; A.N. performed the experiments and interpreted the data; S.K. performed the experiments; R.K. performed the experiments; N.K. performed the experiments and interpreted the data; M.Yamaguchi performed the experiments; S.I. interpreted the data; F.O. interpreted the data; K.N. interpreted the data; H.O. interpreted the data; Y.S. interpreted the data; N.S. performed the experiments and interpreted the data; W.S. interpreted the data; E.K. interpreted the data; J.A. interpreted the data; K.H. interpreted the data; I.K. supervised the project, performed the experiments, interpreted the data, and wrote the manuscript; I.K. had primary responsibility for the final content. All authors read and approved the final version of the manuscript.

## Supplementary information

**Extended Data Fig. 1.**
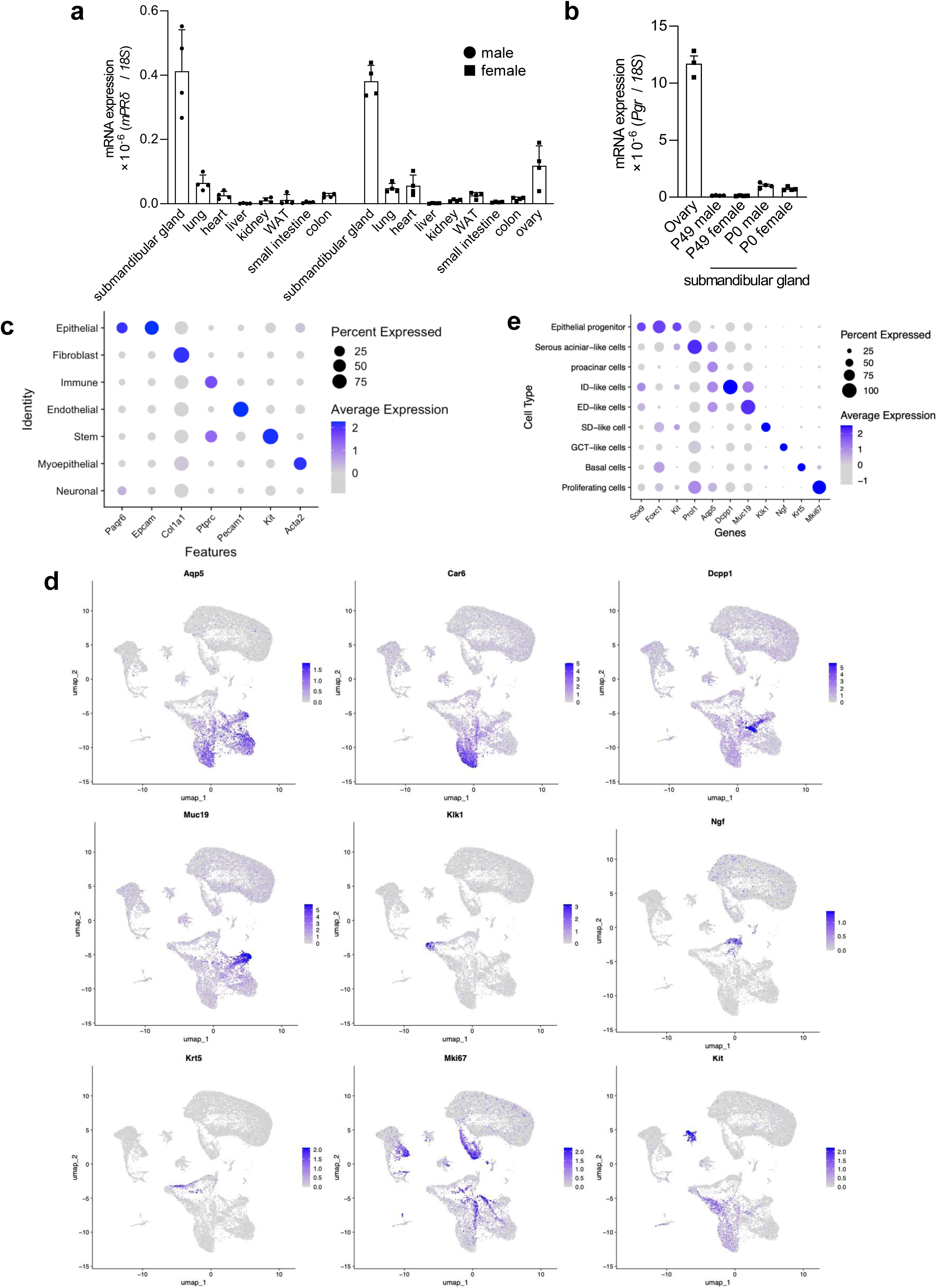
| mPRδ is selectively enriched in the submandibular gland and epithelial compartments. (a) *mPR*δ mRNA expression across tissues from male and female mice at P49, measured by RT-qPCR (*n* = 4). WAT, white adipose tissue (epididymal fat). Expression was normalized to 18S rRNA. (b) Expression of the nuclear progesterone receptor *Pgr* in ovary and SMGs from male and female mice at P0 and P49 (*n* = 3–4), showing minimal expression compared to reproductive tissues. (c) Dot plot showing canonical marker-gene expression across major cell types identified in the P0 SMG single-cell RNA sequencing dataset. Dot size indicates the percentage of expressing cells and color indicates average expression. (d) UMAP visualization of single-cell RNA-seq data from submandibular glands at postnatal day 0 (P0), with feature plots of representative marker genes, including *Aqp5* (proacinar), *Car3* (serous acinar), *Dcpp1* (intercalated duct), *Muc19* (excretory duct), *Klk1* (GCT/striated duct), *Ngf* (GCT), *Krt5* (basal/progenitor), *Mki67* (proliferating), and *Kit* (progenitor). (e) Dot plot showing marker-gene expression across epithelial progenitor, serous acinar-like, proacinar, intercalated duct (ID)-like, excretory duct (ED)-like, striated duct (SD)-like, GCT-like, basal, and proliferating cell populations. Dot size indicates the percentage of expressing cells and color indicates average expression.

**Extended Data Fig. 2.**
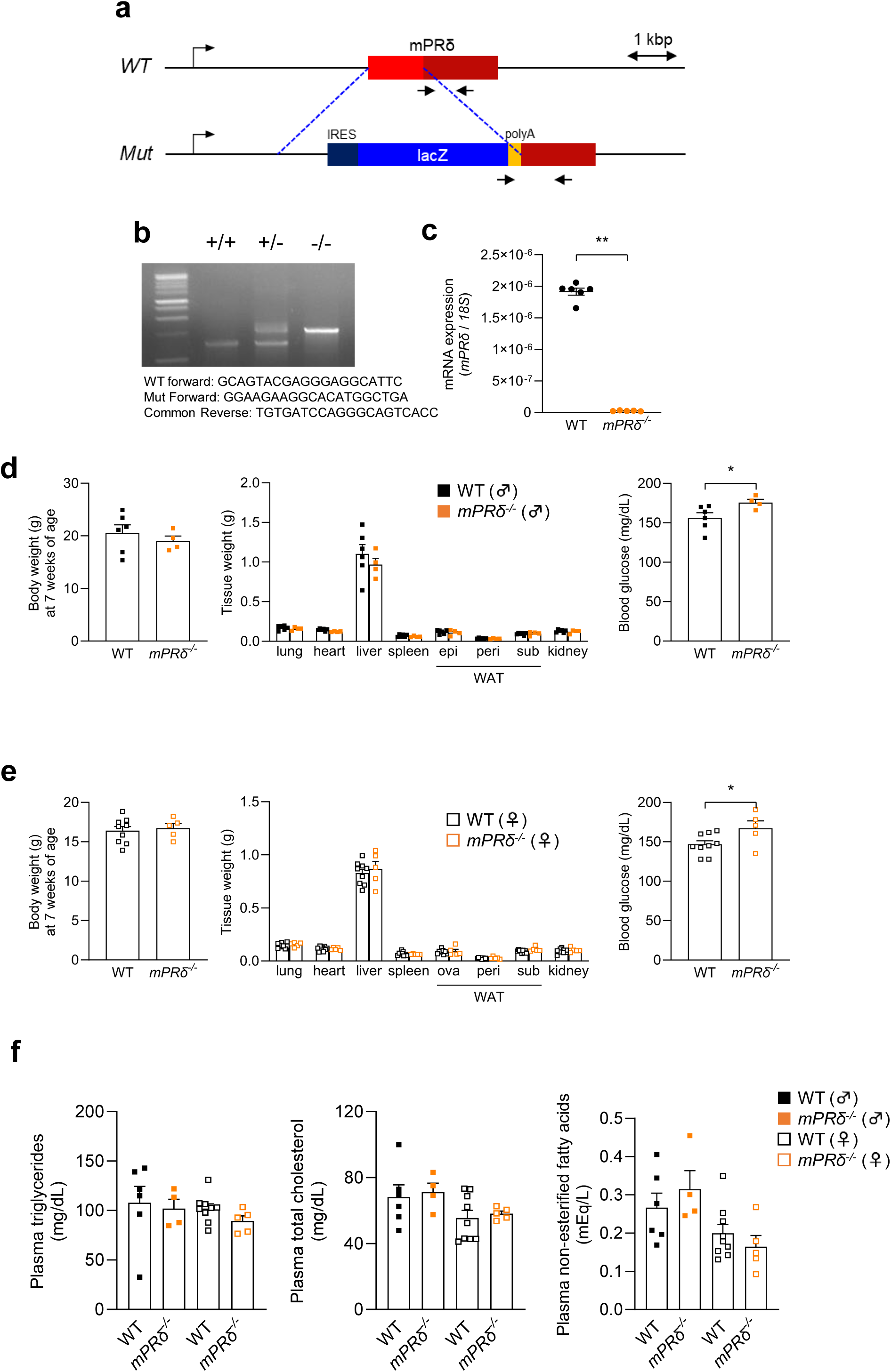
| Generation and validation of mPRδ-deficient mice. (a) Schematic of the targeting strategy for the mPRδ locus. A 1.1-kb genomic fragment containing the exon encoding mPRδ was replaced with a LacZ cassette using homologous recombination in C57BL/6J embryonic stem cells. (b) Genotyping strategy. PCR using primers P1, P2, and P3 distinguishes wild-type (1.2 kb; P1/P3) and mutant (1.7 kb; P2/P3) alleles. (c) *mPR*δ mRNA expression in SMGs from 7-week-old wild-type and *mPR*δ^□/□^ mice, measured by RT-qPCR (*n* = 5–6). Expression was normalized to 18S rRNA. (d, e) Body weight and tissue weights of 7-week-old wild-type and *mPR*δ^□/□^ mice in males (d) and females (e), showing no major differences across tissues. Blood glucose levels are indicated on the right. (*n* = 4–9). (f) Plasma triglycerides, total cholesterol, and non-esterified fatty acids in wild-type and *mPR*δ^□/□^ mice (*n* = 4–9), showing no major abnormalities in plasma lipid parameters. Data are presented as mean ± SEM. Statistical significance was determined using a two-tailed Student’s *t*-test or two-way ANOVA followed by the appropriate post hoc tests, as indicated (\**P* < 0.05, \*\**P* < 0.01).

**Extended Data Fig. 3.**
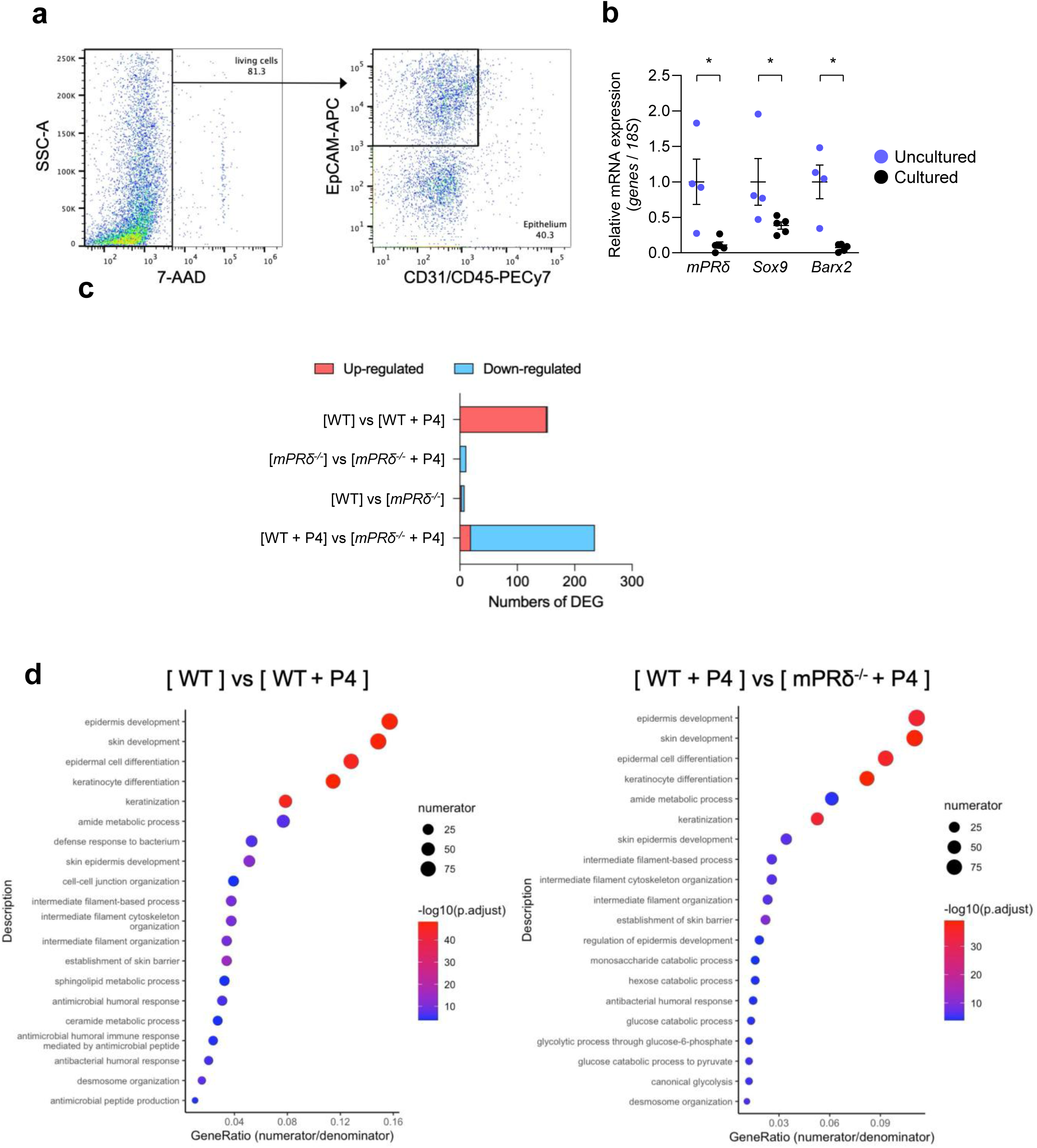
| Progesterone–mPRδ signaling drives epithelial differentiation programs in submandibular gland organoids. (a) Flow-cytometry gating strategy used to isolate epithelial cells from E18 SMGs. Live cells were identified as 7-AAD□, and epithelial cells were defined as CD31□CD45□EpCAM□. (b) mRNA expression of *mPR*δ and epithelial progenitor markers (*Sox9* and *Barx2*) in freshly isolated (uncultured) cells and organoids after culture, measured by RT-qPCR (*n* = 4–5). Expression was normalized to 18S rRNA. (c) RNA sequencing–based differential gene expression analysis of organoids derived from wild-type and *mPR*δ^□/□^ embryos treated with vehicle or P4 (*n* = 3), showing progesterone (P4)-dependent transcriptional responses that are attenuated in *mPR*δ^□/□^ organoids. (d) GO enrichment analysis of differentially expressed genes. Left, vehicle versus P4-treated wild-type organoids; right, P4-treated wild-type versus *mPR*δ^□/□^ organoids. Enriched terms include epithelial differentiation, keratinization, and epidermal development pathways. Adjusted *P* values were calculated using false discovery rate (FDR). Data are presented as mean ± SEM. Statistical significance was determined using a two-tailed Mann–Whitney U test (\**P* < 0.05).

**Extended Data Fig. 4.**
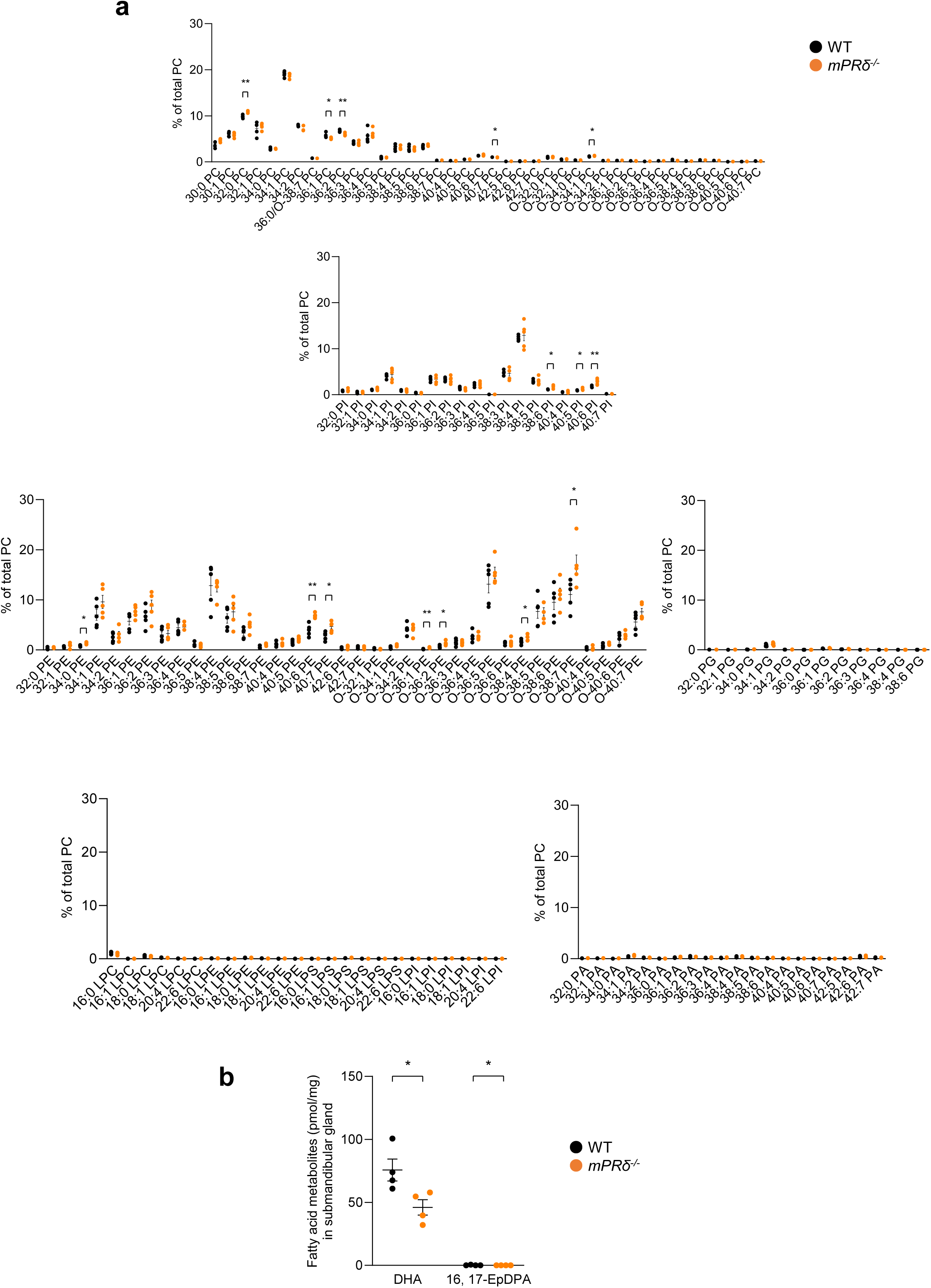
| Membrane phospholipid composition and DHA-derived lipid mediators in mPRδ-deficient submandibular glands. (a) Quantification of membrane phospholipid species across major lipid classes, including phosphatidylcholine (PC), phosphatidylinositol (PI), phosphatidylethanolamine (PE), phosphatidylglycerol (PG), lysophospholipid, and phosphatidic acid (PA), in SMGs from wild-type and *mPR*δ^□/□^ mice at P0 (*n* = 5). Values are expressed as a percentage of total PC content to enable semi-quantitative comparison across phospholipid classes. Several phospholipid species differed significantly between genotypes; among these, PS 40:6 and ether-linked PE O-38:7 were relatively abundant and showed particularly pronounced increases in mPRδ-deficient mice. PS species, including PS 40:6, are shown separately in Fig. 2a. (b) Levels of DHA and the DHA-derived epoxy metabolite 16,17-epoxydocosapentaenoic acid (16,17-EpDPA) in SMGs from wild-type and *mPR*δ^□/□^ mice at P0 (*n* = 4). Data are presented as mean ± SEM. Statistical significance was determined using a two-tailed Mann–Whitney U-test (\**P* < 0.05, \*\**P* < 0.01).

**Extended Data Fig. 5.**
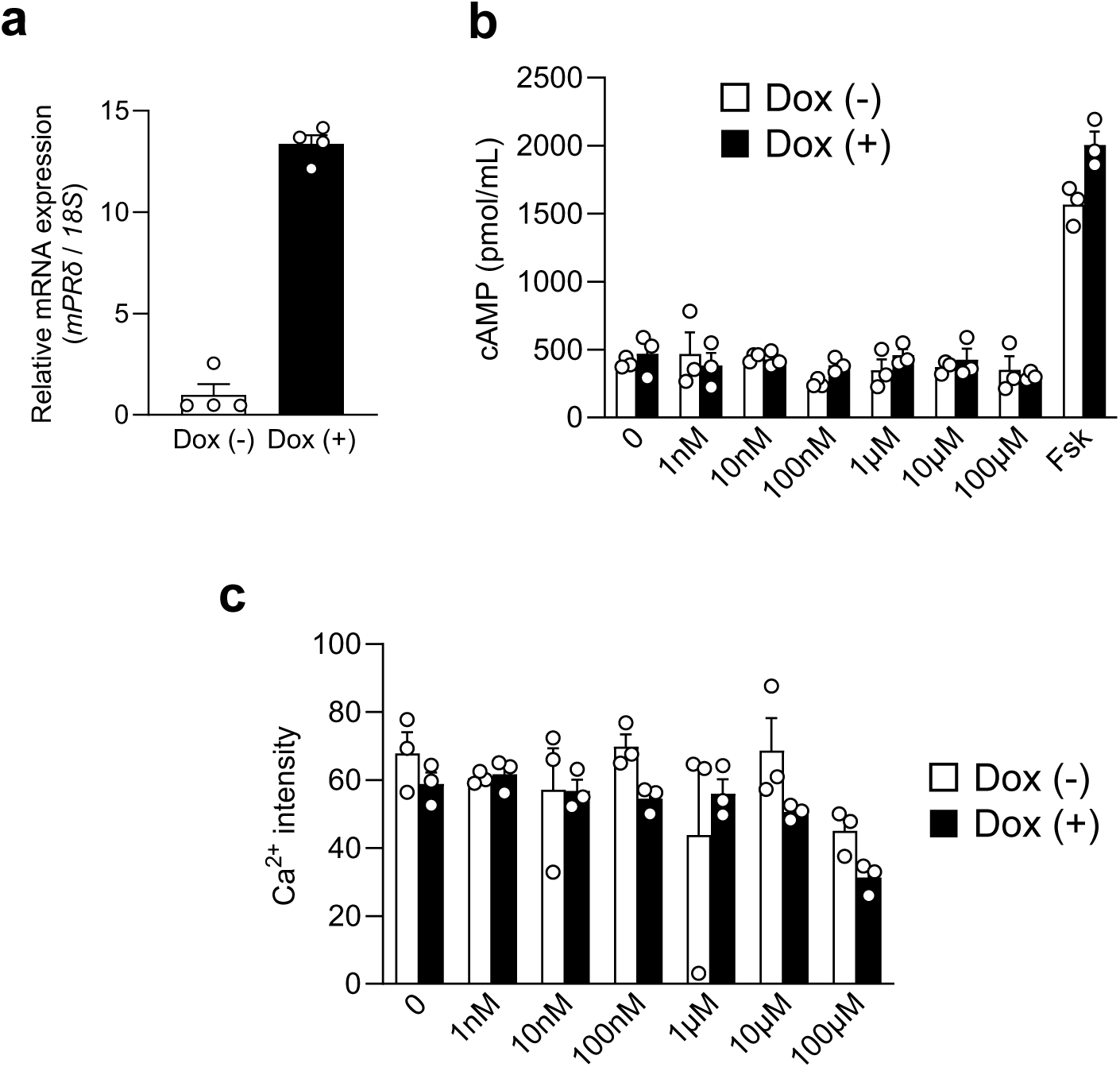
| mPRδ does not engage canonical GPCR signaling pathways in HEK293 cells. (a) *mPR*δ mRNA expression in Flp-In T-REx HEK293 cells with or without doxycycline (Dox; 10 μg/mL for 18 h), measured by RT-qPCR (*n* = 4). Expression was normalized to 18S rRNA. (b) Intracellular cAMP concentrations following stimulation with the indicated concentrations of progesterone in control (Dox□) and mPRδ-expressing (Dox□) cells (*n* = 3). Forskolin (FSK) was used as a positive control. (c) Intracellular Ca^2+^ responses following stimulation with the indicated concentrations of progesterone in Dox^-^ and Dox^+^ cells (*n* = 3). Data are presented as mean ± SEM.

**Extended Data Fig. 6.**
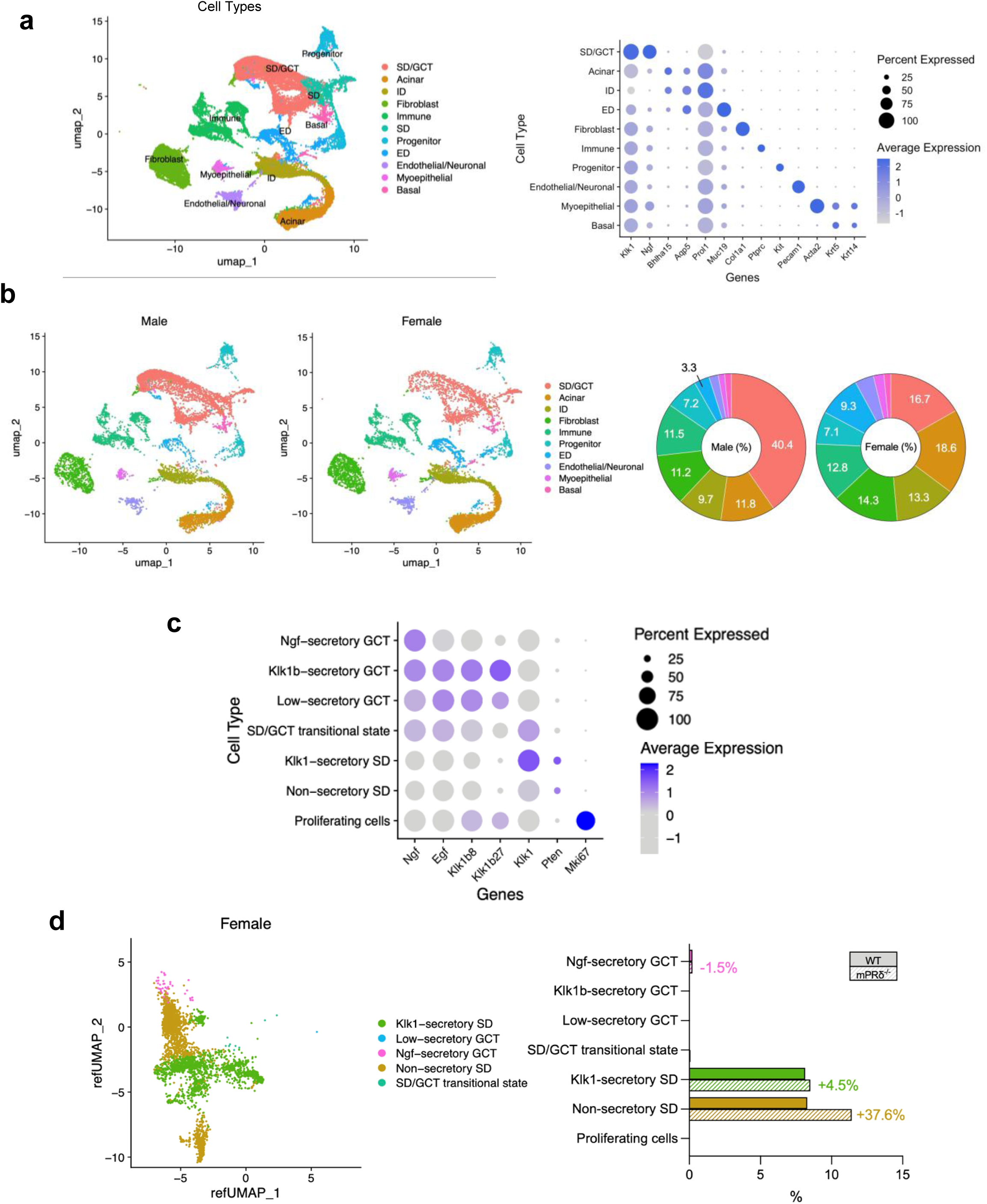
| Single-cell transcriptomic analysis reveals sex- and mPRδ-dependent epithelial composition in adult submandibular glands. (a) UMAP visualization of single-cell RNA sequencing data from SMGs of 7-week-old mice, annotated by major cell type (left). Dot plot showing canonical marker-gene expression across annotated cell types (right). Dot size indicates the percentage of expressing cells and color indicates average expression. (b) UMAP visualizations of SMG cells from male and female mice (left) and donut charts showing the relative proportions of major cell types in male and female SMGs (right). (c) Dot plot showing marker-gene expression across GCT- and SD-associated epithelial subclusters. Dot size indicates the percentage of expressing cells and color indicates average expression. (d) UMAP visualization of epithelial SD/GCT subclusters in female mice (left). Bar plots showing the proportions of each SD/GCT subpopulation in wild-type and *mPR*δ^□/□^ female mice (right).

**Extended Data Fig. 7.**
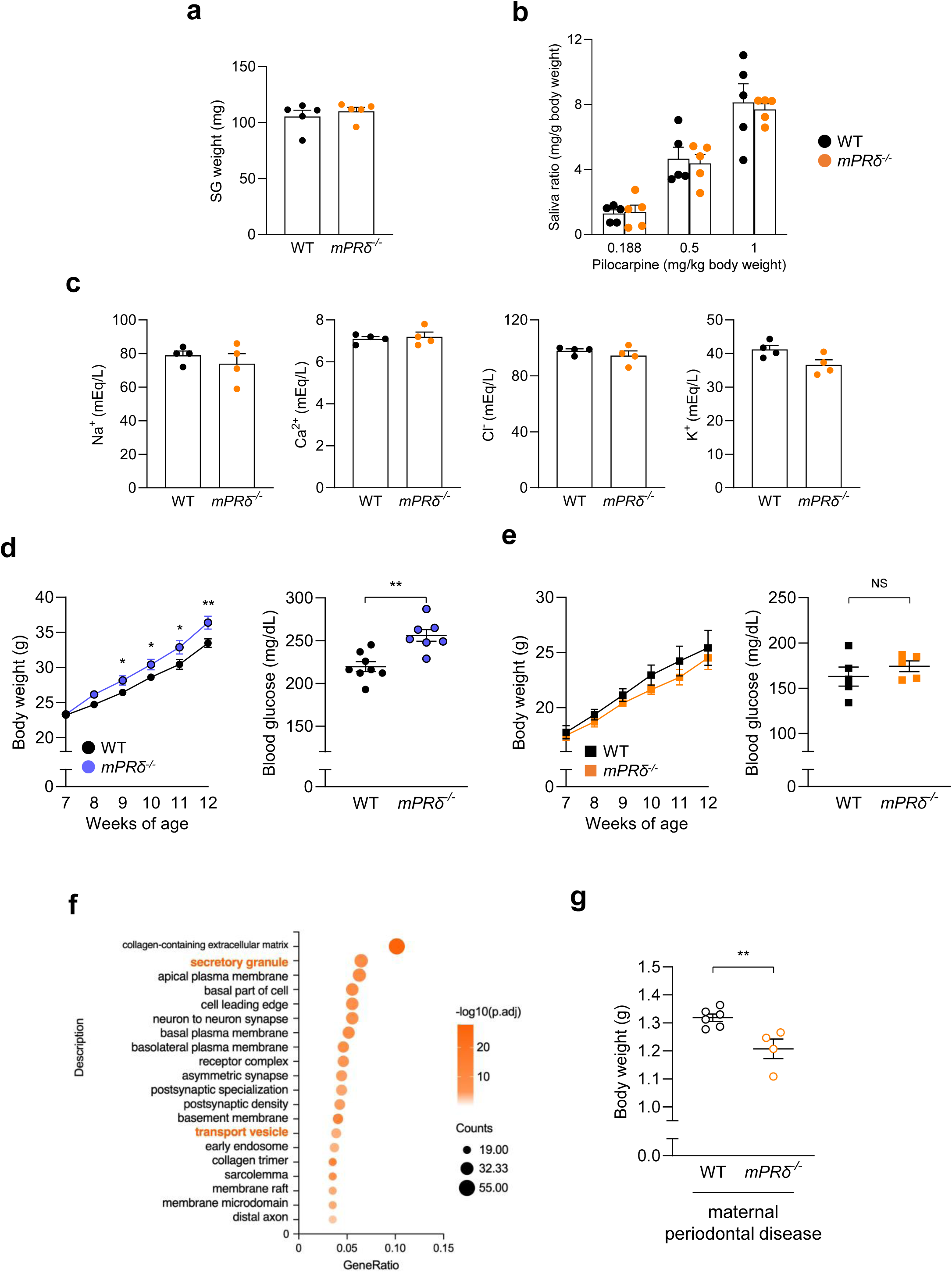
| Sex- and pregnancy-dependent physiological phenotypes associated with mPRδ deficiency. (a) SMG weight in 7-week-old male wild-type and *mPR*δ^□/□^ mice (*n* = 5). (b) Saliva secretion in response to pilocarpine stimulation (0.188–1.0 mg/kg body weight) over 30 minutes in 7-week-old male mice (*n* = 5), showing comparable secretory capacity between genotypes. (c) Electrolyte concentrations in saliva, including Na□, Ca²□, Cl□, and K□, following pilocarpine stimulation in 7-week-old male mice (*n* = 4), indicating preserved ionic composition. (d) Body weight gain and blood glucose levels in wild-type and *mPR*δ^□/□^ male mice during high-fat diet (HFD) feeding from 7 to 12 weeks of age (*n* = 8, 7). *mPR*δ^□/□^ male mice exhibit increased body weight and elevated blood glucose levels compared to wild-type controls. (e) Body weight gain and blood glucose levels in wild-type and *mPR*δ^□/□^ female mice under HFD feeding (*n* = 5), showing no marked differences between genotypes. (f) GO enrichment analysis of differentially expressed genes in SMGs from GD17 wild-type and *mPR*δ^□/□^ mice (*n* = 4). Enriched categories include secretory granule and transport vesicle. (g) Body weight of pups born to wild-type and *mPR*δ^□/□^ mothers following maternal periodontal disease induction (*n* = 6, 4). Data are presented as mean ± SEM. Statistical significance was determined using a two-tailed Mann–Whitney U-test (\**P* < 0.05, \*\**P* < 0.01; NS, not significant).

**Extended Data Fig. 8.**
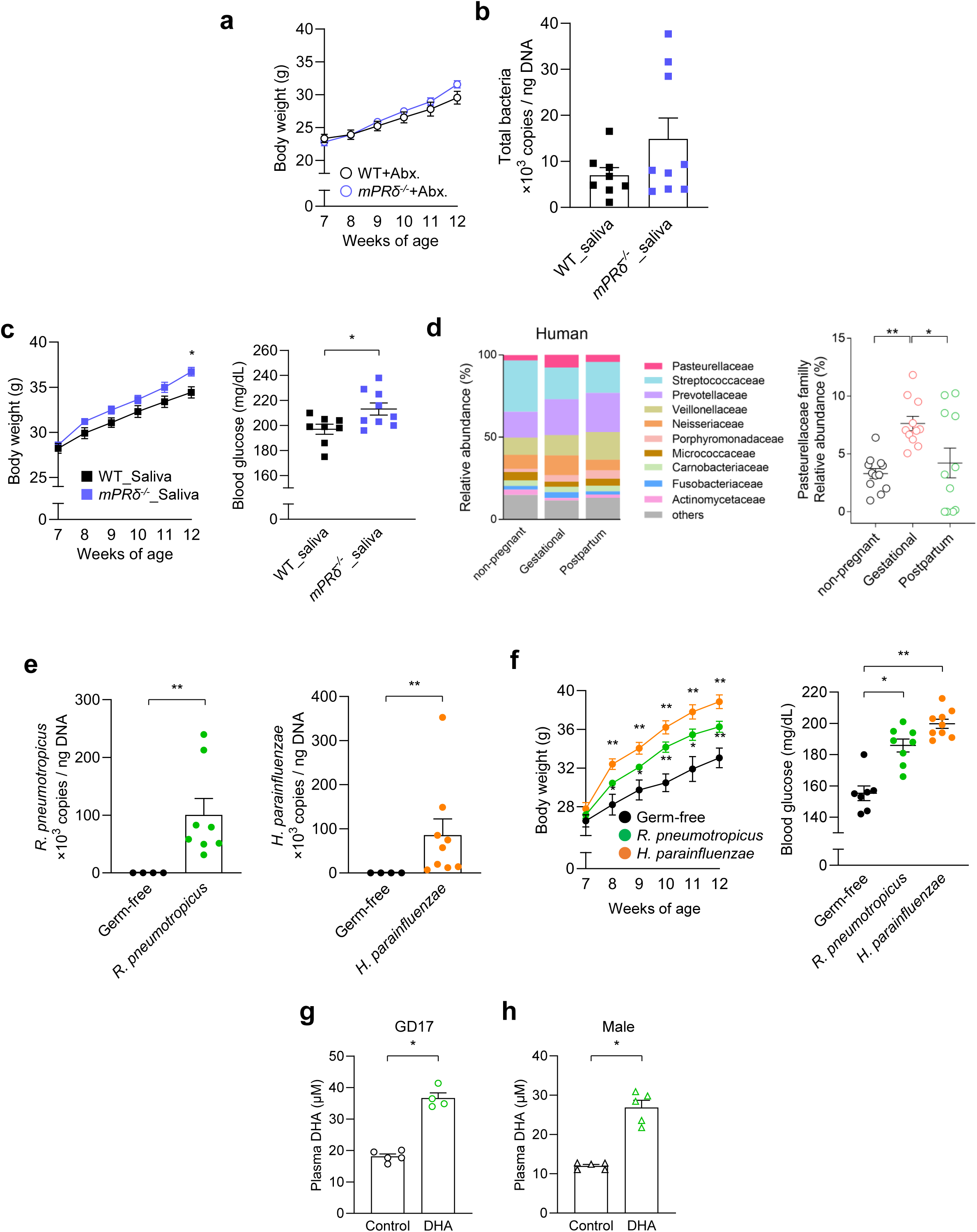
| Oral microbiota and DHA signaling mediate mPRδ deficiency-associated periodontal and metabolic phenotypes. (a) Effects of antibiotic treatment on high-fat diet (HFD)-induced body-weight gain in male wild-type and *mPR*δ^□/□^ mice (*n* = 7–9). (b) qPCR verification of oral microbiota transfer from wild-type or *mPR*δ□/□ donor mice into germ-free recipients (*n* = 8, 9). (c) Metabolic phenotypes in mice colonized with oral microbiota from wild-type or *mPR*δ□/□ donors under HFD conditions (*n* = 8, 9). (d) Relative abundances of major bacterial families in human saliva during non-pregnancy, pregnancy, and the postpartum period (left), and relative abundance of Pasteurellaceae across these stages (right) (*n* = 11–12). (e) qPCR verification of colonization of germ-free mice with *R. pneumotropicus* or *H. parainfluenzae* (*n* = 4–9). (f) Metabolic phenotypes in mice colonized with *R. pneumotropicus* or *H. parainfluenzae* under HFD conditions (*n* = 4–9). (g) Plasma free DHA concentrations in GD17 pregnant *mPR*δ^□/□^ mice with or without DHA supplementation (*n* = 4–5). (h) Plasma free DHA concentrations in 7-week-old male *mPR*δ□/□ mice with or without DHA supplementation (*n* = 5). Data are presented as mean ± SEM. Statistical significance was determined using a two-tailed Mann–Whitney U-test, except for paired human gestational and postpartum samples, which were analyzed using a two-tailed Wilcoxon signed-rank test (\**P* < 0.05, \*\**P* < 0.01).

## Supplementary tables

**Supplementary Table 1.** Composition of experimental diets used for DHA supplementation. Composition of the control and DHA-supplemented diets based on the AIN-93G formulation. DHA was incorporated by replacing an equivalent amount of dietary fat.

| Ingredient, g | AIN93G | AIN93G+DHA |
| --- | --- | --- |
| Casein | 200 | 198 |
| L-Cystine | 3 | 2.97 |
| Corn Starch | 397.486 | 393.51114 |
| Maltodextrin | 132 | 130.68 |
| Sucrose | 100 | 99 |
| Cellulose, BW200 | 50 | 49.5 |
| Soybean Oil | 70 | 69.3 |
| DHA | 0 | 10 |
| t-butylhydroquinone | 0.014 | 0.01386 |
| Mineral Mix S10022G | 35 | 34.65 |
| Vitamin Mix V10037 | 10 | 9.9 |
| Choline Bitartrate | 2.5 | 2.475 |
| Total weight | 1000 | 1000 |
| Energy, kcal/g |  |  |
| % Energy | 4.0 | 4.0 |
| Fat | 20 | 20 |
| Carbohydrate | 64 | 64 |
| Protein | 16 | 16 |

**Supplementary Table 2.**
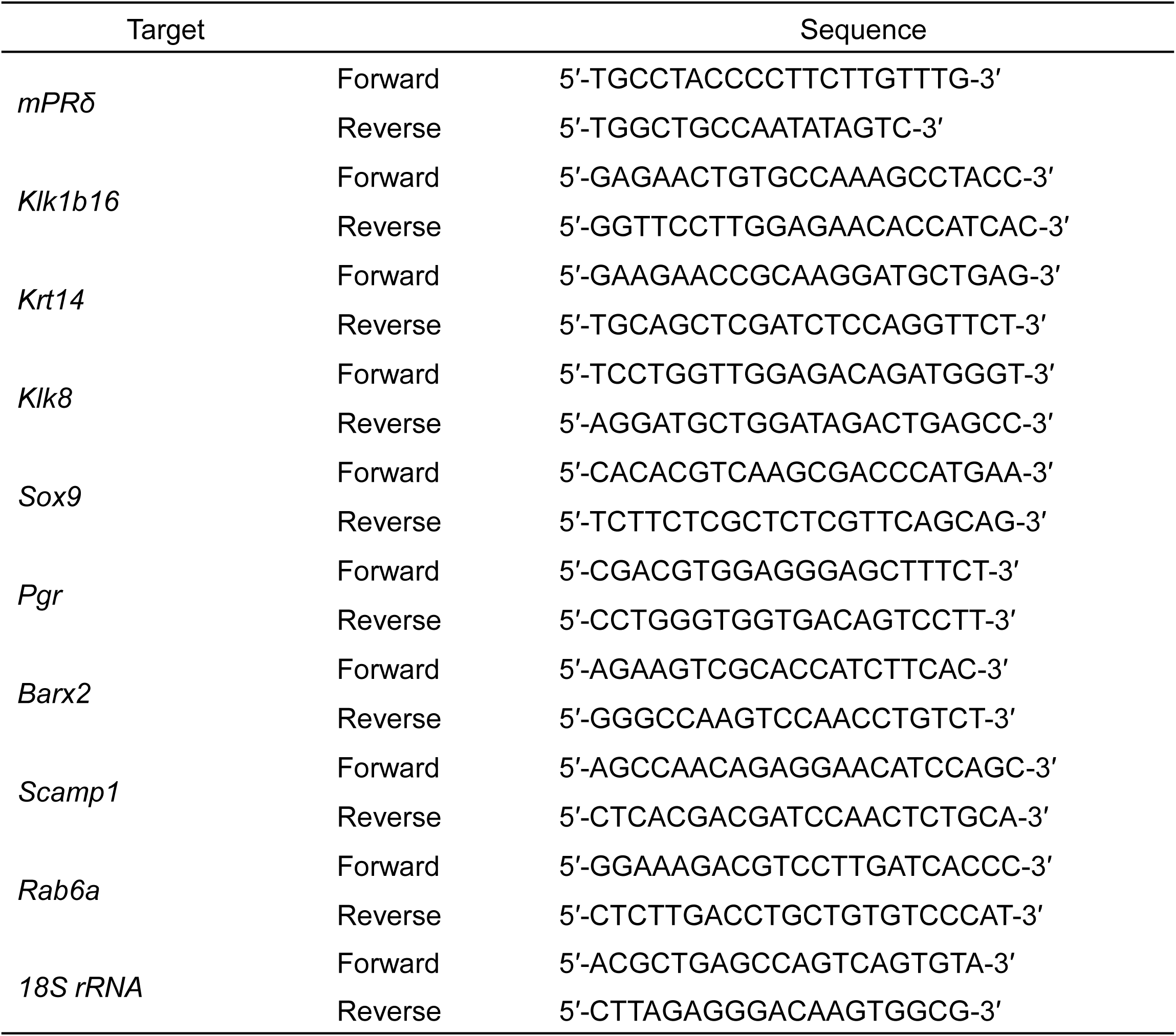
Primer sequences used in this study. Primer sequences used for RT-qPCR and genotyping analyses.

**Supplementary Table 3.**
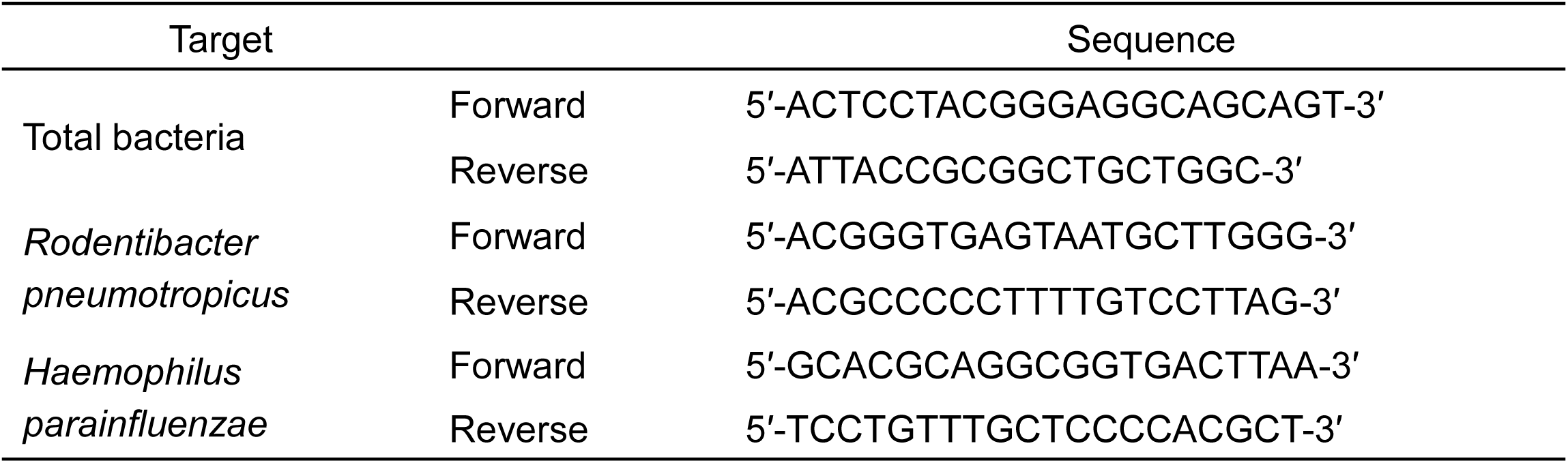
Bacterial primer sequences used in this study. Primer sequences used for qPCR.

## References

1. Baker, J. L., Mark Welch, J. L., Kauffman, K. M., McLean, J. S. & He, X. The oral microbiome: diversity, biogeography and human health. Nat. Rev. Microbiol. 22, 89–104 (2024). 10.1038/s41579-023-00963-6, PubMed: 37700024.

2. Kunath, B. J., De Rudder, C., Laczny, C. C., Letellier, E. & Wilmes, P. The oral–gut microbiome axis in health and disease. Nat. Rev. Microbiol. 22, 791–805 (2024). 10.1038/s41579-024-01075-5, PubMed: 39039286.

3. Tuganbaev, T., Yoshida, K. & Honda, K. The effects of oral microbiota on health. Science 376, 934–936 (2022). 10.1126/science.abn1890, PubMed: 35617380.

4. Freire, M., Nelson, K. E. & Edlund, A. The oral host–microbial interactome: an ecological chronometer of health? Trends Microbiol. 29, 551–561 (2021). 10.1016/j.tim.2020.11.004, PubMed: 33279381.

5. Hajishengallis, G. & Chavakis, T. Local and systemic mechanisms linking periodontal disease and inflammatory comorbidities. Nat. Rev. Immunol. 21, 426–440 (2021). 10.1038/s41577-020-00488-6, PubMed: 33510490.

6. Tonelli, A., Lumngwena, E. N. & Ntusi, N. A. B. The oral microbiome in the pathophysiology of cardiovascular disease. Nat. Rev. Cardiol. 20, 386–403 (2023). 10.1038/s41569-022-00825-3, PubMed: 36624275.

7. Vergnes, J. N. & Sixou, M. Preterm low birth weight and maternal periodontal status: a meta-analysis. Am. J. Obstet. Gynecol. 196, 135.e1–135.e7 (2007). 10.1016/j.ajog.2006.09.028, PubMed: 17306654.

8. Boyapati, R., Cherukuri, S. A., Bodduru, R. & Kiranmaye, A. Influence of female sex hormones in different stages of women on periodontium. J. Midlife Health 12, 263–266 (2021). 10.4103/jmh.jmh_142_21, PubMed: 35264831.

9. Saadaoui, M., Singh, P. & Al Khodor, S. A. Oral microbiome and pregnancy: a bidirectional relationship. J. Reprod. Immunol. 145, 103293 (2021). 10.1016/j.jri.2021.103293, PubMed: 33676065.

10. Nuriel-Ohayon, M., Neuman, H. & Koren, O. Microbial changes during pregnancy, birth, and infancy. Front. Microbiol. 7, 1031 (2016). 10.3389/fmicb.2016.01031, PubMed: 27471494.

11. Chibly, A. M., Aure, M. H., Patel, V. N. & Hoffman, M. P. Salivary gland function, development, and regeneration. Physiol. Rev. 102, 1495–1552 (2022). 10.1152/physrev.00015.2021, PubMed: 35343828.

12. Shang, Y. F. et al. Progress in salivary glands: endocrine glands with immune functions. Front. Endocrinol. 14, 1061235 (2023). 10.3389/fendo.2023.1061235, PubMed: 36817607.

13. Ogawa, M. et al. Functional salivary gland regeneration by transplantation of a bioengineered organ germ. Nat. Commun. 4, 2498 (2013). 10.1038/ncomms3498, PubMed: 24084982.

14. Aure, M. H. et al. FGFR2 is essential for salivary gland duct homeostasis and MAPK-dependent seromucous acinar cell differentiation. Nat. Commun. 14, 6485 (2023). 10.1038/s41467-023-42243-0, PubMed: 37838739.

15. Aure, M. H., Symonds, J. M., Mays, J. W. & Hoffman, M. P. Epithelial cell lineage and signaling in murine salivary glands. J. Dent. Res. 98, 1186–1194 (2019). 10.1177/0022034519864592, PubMed: 31331226.

16. Saunders, F. J. Effects of sex steroids and related compounds on pregnancy and on development of the young. Physiol. Rev. 48, 601–643 (1968). 10.1152/physrev.1968.48.3.601, PubMed: 4873867.

17. López-García, C. et al. Molecular and morphological changes in placenta and embryo development associated with the inhibition of polyamine synthesis during midpregnancy in mice. Endocrinology 149, 5012–5023 (2008). 10.1210/en.2008-0084, PubMed: 18583422.

18. Dey, A. K. et al. Salivary proteome signatures in the early and middle stages of human pregnancy with term birth outcome. Sci. Rep. 10, 8022 (2020). 10.1038/s41598-020-64483-6, PubMed: 32415095.

19. Lösel, R. M. et al. Nongenomic steroid action: controversies, questions, and answers. Physiol. Rev. 83, 965–1016 (2003). 10.1152/physrev.00003.2003, PubMed: 12843413.

20. Garg, D., Ng, S. S. M., Baig, K. M., Driggers, P. & Segars, J. Progesterone-mediated non-classical signaling. Trends Endocrinol. Metab. 28, 656–668 (2017). 10.1016/j.tem.2017.05.006, PubMed: 28651856.

21. Thomas, P. Characteristics of membrane progestin receptor alpha (mPRalpha) and progesterone membrane receptor component 1 (PGRMC1) and their roles in mediating rapid progestin actions. Front. Neuroendocrinol. 29, 292–312 (2008). 10.1016/j.yfrne.2008.01.001, PubMed: 18343488.

22. Kasubuchi, M. et al. Membrane progesterone receptor beta (mPRβ/Paqr8) promotes progesterone-dependent neurite outgrowth in PC12 neuronal cells via non-G protein-coupled receptor (GPCR) signaling. Sci. Rep. 7, 5168 (2017). 10.1038/s41598-017-05423-9, PubMed: 28701790.

23. Smith, J. L. et al. Heterologous expression of human mPRalpha, mPRbeta and mPRgamma in yeast confirms their ability to function as membrane progesterone receptors. Steroids 73, 1160–1173 (2008). 10.1016/j.steroids.2008.05.003, PubMed: 18603275.

24. Nader, N. et al. Progesterone induces meiosis through two obligate co-receptors with PLA2 activity. eLife 13, RP92635 (2025). 10.7554/eLife.92635, PubMed: 39873665.

25. Watanabe, K. et al. Maternal progesterone and adipose mPRε in pregnancy regulate the embryonic nutritional state. Cell Rep. 44, 115433 (2025). 10.1016/j.celrep.2025.115433, PubMed: 40085645.

26. Zou, T. et al. PAQR6 as a prognostic biomarker and potential therapeutic target in kidney renal clear cell carcinoma. Front. Immunol. 15, 1521629 (2024). 10.3389/fimmu.2024.1521629, PubMed: 39742277.

27. Kimura, I. et al. Maternal gut microbiota in pregnancy influences offspring metabolic phenotype in mice. Science 367, eaaw8429 (2020). 10.1126/science.aaw8429, PubMed: 32108090.

28. Chatzeli, L., Gaete, M. & Tucker, A. S. Fgf10 and Sox9 are essential for the establishment of distal progenitor cells during mouse salivary gland development. Development 144, 2294–2305 (2017). 10.1242/dev.146019, PubMed: 28506998.

29. Naka, T. & Yokose, S. Immunohistochemical localization of barx2 in the developing fetal mouse submandibular glands. Acta Histochem. Cytochem. 42, 47–53 (2009). 10.1267/ahc.08027, PubMed: 19492027.

30. Hauser, B. R. et al. Generation of a single-cell RNAseq atlas of murine salivary gland development. iScience 23, 101838 (2020). 10.1016/j.isci.2020.101838, PubMed: 33305192.

31. Gleeson, P. J., Camara, N. O. S., Launay, P., Lehuen, A. & Monteiro, R. C. Immunoglobulin A antibodies: from protection to harmful roles. Immunol. Rev. 328, 171–191 (2024). 10.1111/imr.13424, PubMed: 39578936.

32. Szczuko, M. et al. The role of arachidonic and linoleic acid derivatives in pathological pregnancies and the human reproduction process. Int. J. Mol. Sci. 21, 9628 (2020). 10.3390/ijms21249628, PubMed: 33348841.

33. Takagaki, T., Shingaki, T., Kono, N. & Aoki, J. Emerging roles of CREST superfamily members in lipid metabolism. J. Lipid Res. 101117 (2026). 10.1016/j.jlr.2026.101117, PubMed: 42551530.

34. Nakamura, M. T., Yudell, B. E. & Loor, J. J. Regulation of energy metabolism by long-chain fatty acids. Prog. Lipid Res. 53, 124–144 (2014). 10.1016/j.plipres.2013.12.001, PubMed: 24362249.

35. Kimura, I., Ichimura, A., Ohue-Kitano, R. & Igarashi, M. Free fatty acid receptors in health and disease. Physiol. Rev. 100, 171–210 (2020). 10.1152/physrev.00041.2018, PubMed: 31487233.

36. Oh, D. Y. et al. GPR120 is an omega-3 fatty acid receptor mediating potent anti-inflammatory and insulin-sensitizing effects. Cell 142, 687–698 (2010). 10.1016/j.cell.2010.07.041, PubMed: 20813258.

37. Miyamoto, J. et al. Gut microbiota confers host resistance to obesity by metabolizing dietary polyunsaturated fatty acids. Nat. Commun. 10, 4007 (2019). 10.1038/s41467-019-11978-0, PubMed: 31488836.

38. Gao, S. et al. Oral microbiome modulation mitigates hyperglycemia exacerbation in gestational diabetes mellitus. Nat. Commun. (2026). 10.1038/s41467-026-74917-w, PubMed: 42399225.

39. Darveau, R. P. Periodontitis: a polymicrobial disruption of host homeostasis. Nat. Rev. Microbiol. 8, 481–490 (2010). 10.1038/nrmicro2337, PubMed: 20514045.

40. Bagavant, H. et al. A method for the measurement of salivary gland function in mice. J. Vis. Exp. 25, 57203 (2018). 10.3791/57203, PubMed: 29443033.

41. Ikeda, T. et al. The free fatty acid receptor GPR164 maintains intestinal homeostasis and barrier function. EMBO Rep. 26, 5905–5930 (2025). 10.1038/s44319-025-00611-5, PubMed: 41152604.

42. Callahan, B. J. et al. DADA2: high-resolution sample inference from Illumina amplicon data. Nat. Methods 13, 581–583 (2016). 10.1038/nmeth.3869, PubMed: 27214047.

43. Callahan, B. J. et al. High-throughput amplicon sequencing of the full-length 16S rRNA gene with single-nucleotide resolution. Nucleic Acids Res. 47, e103 (2019). 10.1093/nar/gkz569, PubMed: 31269198.

44. Shimizu, H. et al. Sucrose-preferring gut microbes prevent host obesity by producing exopolysaccharides. Nat. Commun. 16, 1145 (2025). 10.1038/s41467-025-56470-0, PubMed: 39880823.

45. Abe, T. & Hajishengallis, G. Optimization of the ligature-induced periodontitis model in mice. J. Immunol. Methods 394, 49–54 (2013). 10.1016/j.jim.2013.05.002, PubMed: 23672778.

46. Li, Y. et al. Salivary mycobiome dysbiosis and its potential impact on bacteriome shifts and host immunity in oral lichen planus. Int. J. Oral Sci. 11, 13 (2019). 10.1038/s41368-019-0045-2, PubMed: 31263096.

